# A Human Single-cell Atlas Identifies OLR1+ Scar-associated Macrophages as a Therapeutic Target for Chronic Liver Disease

**DOI:** 10.1101/2025.08.27.671899

**Authors:** Eleni Papachristoforou, Kexin Kong, Fabio Colella, Jacky Tam, Juliet Luft, Elena F. Sutherland, Stefan Veizades, Malgorzata Grzelka, Pak Kwan Qiu, Ayma Asif, Ines Battle, Max Hammer, John R. Wilson-Kanamori, George Finney, Ginerva Pistocchi, Alvile Kasarinaite, Gareth-Rhys Jones, Calum C. Bain, Laura J. Pallett, Catalina A. Vallejos, Neil C. Henderson, David C. Hay, Timothy J Kendall, Jonathan A Fallowfield, Prakash Ramachandran

**Affiliations:** Centre for Inflammation Research, Institute for Regeneration and Repair, University of Edinburgh, Edinburgh, United Kingdom; Centre for Cardiovascular Science, University of Edinburgh, UK; Division of Infection and Immunity, Institute of Immunity and Transplantation, University College London, London, United Kingdom; Centre for Regenerative Medicine, Institute for Regeneration and Repair, University of Edinburgh, Edinburgh, United Kingdom; MRC Institute of Genetics and Cancer, University of Edinburgh, Edinburgh, United Kingdom; The Alan Turing Institute, London, United Kingdom; MRC Human Genetics Unit, Institute of Genetics and Cancer, University of Edinburgh, Edinburgh, United Kingdom; Edinburgh Pathology, University of Edinburgh, Edinburgh, United Kingdom

## Abstract

Chronic liver disease (CLD) is a major global healthcare problem. Irrespective of cause, chronic damage to the liver results in fibrosis, which is associated with adverse clinical outcomes. Immune cells, in particular monocyte-derived macrophages (MDMs), are key regulators of fibrosis and represent an attractive therapeutic target for CLD. However, it has remained unclear which specific subpopulation of MDMs drives pro-inflammatory and pro-fibrotic functions in human CLD and how they might be selectively modulated. Here, we generate an annotated human liver single cell atlas from 42 healthy and 35 CLD patients, identifying 125 transcriptionally distinct cellular states. Leveraging large patient and cell numbers, our atlas resolves rare liver cell states and distinguishes two types of disease-expanded scar-associated macrophages (SAMac), including a specific subpopulation with a pro-inflammatory pro-fibrotic phenotype. The scavenger receptor OLR1 was enriched in pro-inflammatory SAMacs and high hepatic OLR1 expression was associated with increased mortality in CLD patients. A corresponding monocyte-derived OLR1+ SAMac subpopulation expanded in a mouse model of CLD and exhibited a pro-inflammatory phenotype based on single-cell RNA-seq, single-cell ATAC-seq, and flow cytometry analyses. Primary human OLR1+ MDMs promoted fibrogenic signalling in multilineage liver spheroid cultures, whilst specific targeting of OLR1 reduced IL-1β production by human macrophages and attenuated myofibroblast activation. Overall, our annotated human liver single-cell atlas provides a valuable reference to study disease-associated cell states in CLD. We utilise this resource to identify a distinct pro-inflammatory subpopulation of SAMacs and highlight OLR1 as a potential therapeutic target to specifically modulate SAMac function and attenuate liver fibrosis.

## Main

Chronic liver disease (CLD) is a major global healthcare problem, affecting 480 million people worldwide and resulting in approximately 2 million deaths per year^1^. Irrespective of cause, chronic damage to the liver results in progressive scarring (fibrosis) which can ultimately advance to cirrhosis, leading to clinical complications and increased mortality. Higher levels of fibrosis in the liver are associated with poor clinical outcomes^2,3^, whereas improvements in fibrosis correlate with a better prognosis^4^. Due to this close relationship between fibrosis and CLD outcomes, there is significant interest in developing new antifibrotic therapies for patients with liver diseases. However, such therapies remain very limited, with only one FDA-approved drug (Resmiterom) licenced for the treatment of metabolic dysfunction-associated steatotic liver disease (MASLD)^5^, the commonest cause of CLD in the developed world. Consequently, there is an urgent need for novel therapeutic strategies for patients with CLD.

As with many complex diseases, the application of single-cell RNA-sequencing (scRNAseq) is transforming our understanding of human CLD pathogenesis and facilitating the identification of novel therapeutic targets^6,7^. However, most existing human liver scRNAseq studies include relatively small numbers of cells from a limited number of patient samples, constraining their ability to detect rare cell types or resolve subtle transcriptional states relevant to CLD pathogenesis. Large-scale single-cell atlases have started to address this issue in other human organs such as breast^8^, lung^9^ and gut^10^, but a comprehensive fully annotated human liver scRNAseq atlas spanning healthy and diseased states remains unavailable.

One of the key pathological features of human CLD is the accumulation of immune cells which are important regulators of tissue injury responses and fibrogenesis. Specifically, cells of the mononuclear phagocyte (MP) lineage have been shown to play a crucial role in regulating fibrosis in the liver^11–13^. Previous work using scRNAseq identified TREM2+ CD9+ scar-associated macrophages (SAMacs), which expand in fibrotic human livers and localise to areas of scarring (the fibrotic niche)^14^. These SAMacs promote activation of the scar-producing hepatic myofibroblasts in the liver^14,15^, making them a potential target for antifibrotic therapies. However, hepatic macrophages also play a crucial role in fibrosis regression and tissue repair^16–19^. Indeed, molecules such as TREM2 and SPP1, which are expressed by SAMacs, possess anti-inflammatory and antifibrotic properties^20–23^. Due to the limited resolution of previous studies, it remains unclear whether distinct SAMac subpopulations mediate divergent functions and how pro-inflammatory pro-fibrotic SAMacs might be selectively targeted without compromising reparative macrophage activity.

Macrophages within the diseased liver are a key source of inflammatory mediators which regulate CLD pathogenesis. Specifically, danger-associated molecular patterns (DAMPs) released in response to tissue injury are sensed by macrophage pattern recognition receptors (e.g. toll like receptors (TLRs) or nucleotide-binding oligomerisation domain-like receptors (NLRs)), resulting in NF-κB pathway activation and transcription of pro-inflammatory genes such as TNF-α, IL-6 and IL-1, as well as inflammasome components such as NLRP3)^24^. Among these, IL-1β has been shown to promote activation^25^ and survival^26^ of liver myofibroblasts. Indeed, the key role of macrophage-derived IL-1β in fibrosis pathogenesis is further supported by studies in human cardiac disease^27,28^, where it regulates pro-fibrotic macrophage-fibroblast cross-talk^29^. However, the identity of the specific macrophage subpopulation responsible for IL-1β expression within the human liver fibrotic niche, and the availability of tractable surface targets for therapeutic modulation, remain poorly defined.

OLR1 (LOX-1) is a class E scavenger receptor, implicated in the pathogenesis of atherosclerosis^30^, cardiac fibrosis^31^ and various cancers ^32^. Indeed, therapeutic targeting of OLR1 has been suggested as a treatment for atherosclerosis^30^ with candidate inhibitors currently being explored in clinical trials in patients with cardiovascular disease^33^. Mechanistically, OLR1 signalling can induce pro-inflammatory responses, including activation of the NF-κB pathway and/or NLRP3 inflammasome in endothelial cells^34,35^ and macrophages^36,37^ in the context of atherosclerosis. However, the expression of OLR1 in liver macrophages has not been reported, and its role in liver inflammation and fibrosis remains unexplored.

Here, we generate a fully annotated human liver scRNAseq atlas comprising 77 healthy and CLD livers. This resource reveals previously unrecognised heterogeneity within the hepatic macrophage compartment and identifies a distinct subpopulation of OLR1^+^ SAMacs which expand in CLD and exhibit pro-inflammatory properties. Importantly, high hepatic expression of OLR1 is predictive of adverse clinical outcomes in patients with CLD. Moreover, selective targeting of OLR1+ macrophages reduces IL-1β production and fibrogenic activity in human *in vitro* models. Our scRNAseq atlas provides a foundational resource to study disease-associated cell states and identify novel therapeutic targets in CLD, highlighting OLR1 inhibition as a promising candidate to mitigate hepatic inflammation and fibrosis.

## Results

### Establishing a human CLD single cell atlas

To enable comprehensive analysis of liver cellular heterogeneity and identify disease-associated subpopulations, we first developed a computational workflow to facilitate integrated analysis of data from 7 previously published human scRNAseq studies (Fig. 1A, Supplementary table 1). In total, data was combined from 77 patients (42 healthy and 35 CLD), derived from 38 males (21 healthy and 17 CLD) and 39 females (21 healthy and 18 CLD) based on sample annotation using expression of Y chromosome genes. Following initial QC and data integration, we generated a lineage annotation overview based on previously defined canonical marker genes (Extended Data Fig. 1A, B). Integration was effective across studies (Extended Data Fig. 1C). Each lineage was then subclustered at high resolution to identify and remove low quality (median nFeatures<800) and doublet clusters (co-expression of multiple lineage marker genes), followed by manual annotation of cell types within each lineage, based on curated marker gene expression and previous literature. Annotated lineage subclusters were then combined to generate the final liver atlas, consisting of 649,295 high quality single cells.

**Figure 1:**
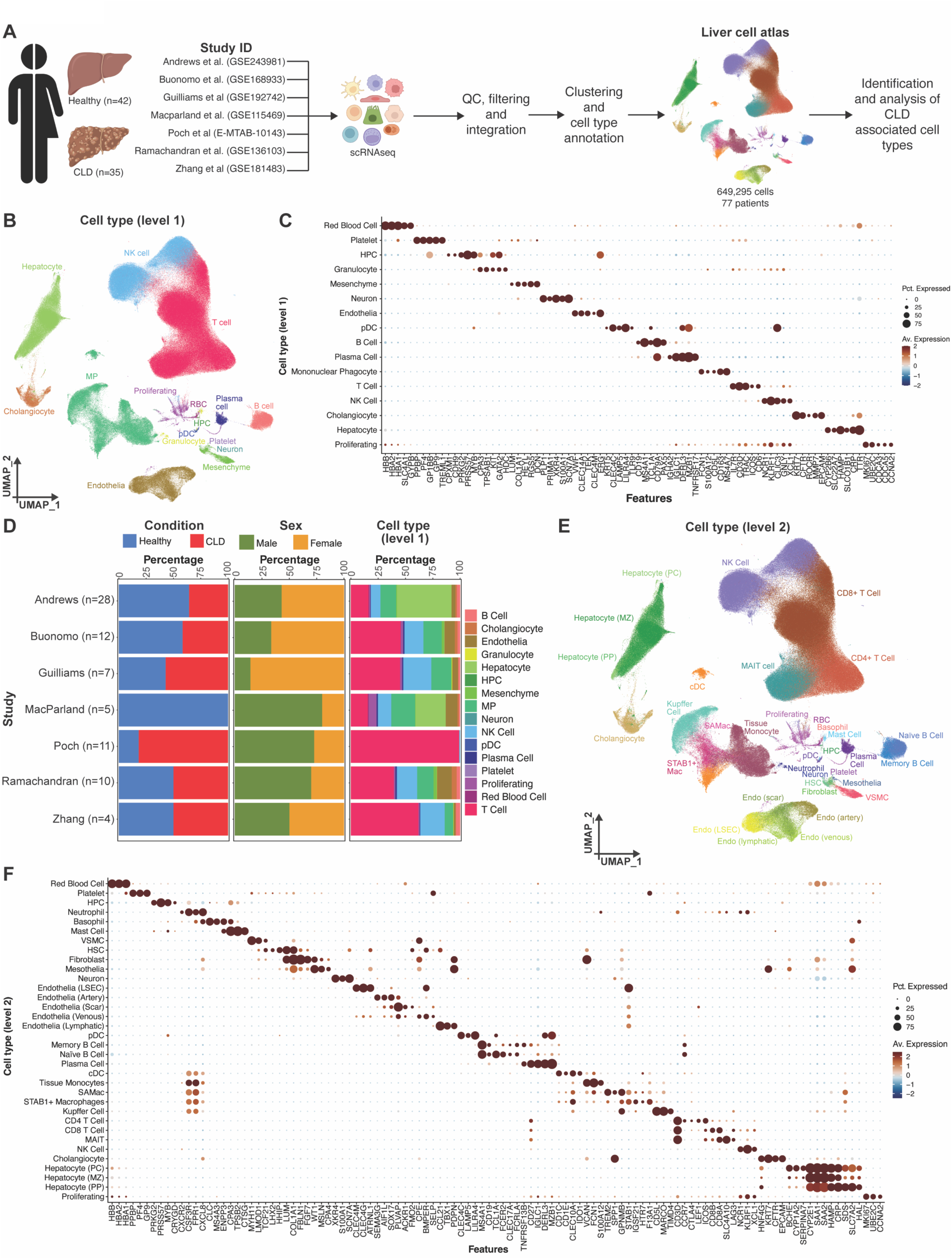
Developing a human liver single-cell atlas. (**A**) Overview of analytical workflow for scRNAseq atlas generation. Created in BioRender. Colella, F. (2025) https://BioRender.com/1ejtusn. (**B**) UMAP of scRNAseq data (649,295 cells; 77 patients) coloured and labelled by level 1 annotation. (**C**) Dotplot showing mean expression of marker genes and percentage of cells expressing them for each level 1 cell type. (**D**) Stacked barplots of proportions of patients of each condition (left; healthy blue, CLD red), patients of each sex (middle; male green, female yellow) and cells of each level 1 cell type (right) across the 7 scRNAseq studies. (**E**) UMAP of scRNAseq data (649,295 cells; 77 patients) coloured and labelled by level 2 annotation. (**F**) Dotplot showing mean expression of marker genes and percentage of cells expressing them for each level 2 cell type.

Three levels of final atlas annotation were established, based on transcriptional similarity between refined clusters (Extended Data Fig. 2A-C). Level 1 clusters consisted of 16 broad cell types (Fig. 1B) with clear distinguishing marker gene profiles (Fig. 1C and Supplementary table 2). All studies contributed to the level 1 annotation, with expected study specific differences based on cell isolation methods from the original publications, such as the inclusion of more epithelial cells in the studies from Andrews et al.^38^ and Macparland et al.^39^ and the inclusion of only T cells from the study by Poch et al.^40^ (Fig. 1D). Apart from the study by Macparland et al., all studies contained a mixture of healthy and CLD patients. All studies contributed both male and female donors to the atlas (Fig. 1D).

Level 2 annotation resulted in 34 cell types (Fig. 1E), with distinct marker gene profiles (Fig. 1F, Supplementary table 3). Level 3 clustering was based on the refined annotation of each lineage subclustering and resulted in a total of 125 final clusters (Extended Data Fig. 2C), with distinct marker gene profiles (Supplementary table 4). The overall atlas structure and hierarchy is summarised in Extended Data Fig. 3. Notably, our atlassing approach from a large number of patients, enabled the identification and annotation of rare cell types such as neurons, mast cells, basophils, neutrophils and haematopoietic progenitor cells (HPCs) (Fig. 1E-F), which have not been well described in individual scRNAseq studies. Fine grained level 3 clusters from each lineage were then used for downstream analysis of changes in specific cell type composition and gene expression between healthy and CLD liver tissue.

To enable user-friendly data browsing of gene expression profiles at the different levels of atlas annotation, we have created a publicly available data browser which is available at: https://shiny.igc.ed.ac.uk/Human_Liver_scRNAseq_Atlas/.

### Identifying scar-associated macrophage heterogeneity in human CLD

To study MP heterogeneity and changes in CLD, we proceeded to analyse the level 3 clustering data. In total 81,577 non-proliferating MPs were included, which partitioned into 14 clusters (Fig. 2A). MP subpopulations were detected from each study included in the CLD atlas, apart from the Poch et al. study which focussed on T cells (Extended Data Fig. 4A, B). MP clusters could broadly be separated into 4 *MNDA*+ *FCN1*+ tissue monocyte subpopulations (TMo1-4), 2 conventional dendritic cell populations (*CD1C*+ *FCER1A*+ cDC2 and migratory *IDO1*+ *CCR7*+ cDC), 3 *CD5L*+ *MARCO*+ Kupffer cell (KC) populations (Periportal (PP) KCs which were high in expression of *IL10* as recently described^41^, *TIMD4*-*PIEZO2*-KC2 and *TIMD4*+ *PIEZO2*+ KC3) and 2 macrophage populations which were high in *STAB1* and *IGSF21* and reminiscent of monocyte-derived KCs (MoKCs) defined in mice^42^. Furthermore, we were able to distinguish 2 distinct populations of macrophages which both expressed high levels of previously defined SAMac marker genes *TREM2*, *CD9, CD63* and *SPP1*^14,15^ (Fig. 2B, Supplementary Table 5). Hence, these subpopulations were annotated SAMac1 and SAMac2.

**Figure 2:**
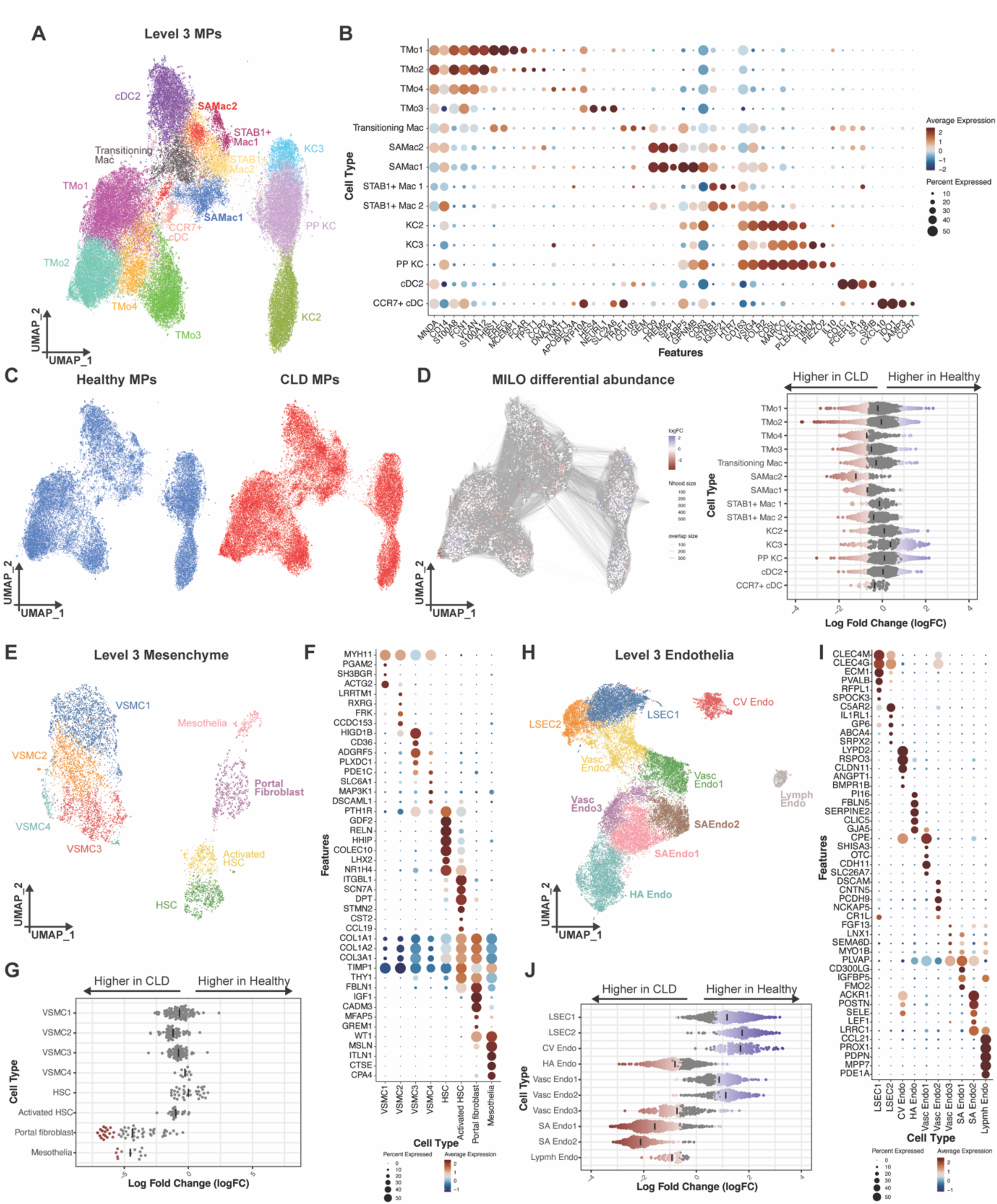
Defining changes in non-parenchymal cell composition in CLD. (**A**) UMAP of mononuclear phagocytes (81,577 cells) coloured and labelled by level 3 annotation. (**B**) Dotplot showing mean expression of marker genes and percentage of cells expressing them for each level 3 MP cell type. (**C**) UMAP of mononuclear phagocytes coloured by patient condition. (**D**) Milo cell neighbourhood differential abundance plots (left) of the significant (FDR <0.05) changes in the MP composition with CLD (colour gradient scale representing log fold changes). Blue represents enrichment in healthy while red denotes expansion in CLD. Beeswarm plot (right) of the log fold changes in the Milo neighbourhoods with CLD status, grouped into each level MP cell-type subcluster. Neighbourhoods with a significant change in cellular abundance are coloured as indicated. Black lines denote mean logFC for each cell type. (**E**) UMAP of mesenchymal cells (7,301 cells) coloured and labelled by level 3 annotation. (**F**) Dotplot showing mean expression of marker genes and percentage of cells expressing them for each level 3 mesenchymal cell type. (**G**) Beeswarm plot of the log fold changes in Milo cell neighbourhoods in mesenchymal cells by CLD status. Milo neighbourhoods grouped into each level 3 mesenchymal cell-type subcluster. Neighbourhoods with a significant (FDR <0.05) change in cellular abundance are coloured with blue representing enrichment in healthy and red denoting expansion in CLD. Black lines denote mean logFC for each cell type. (**H**) UMAP of endothelial cells (36,591 cells) coloured and labelled by level 3 annotation. (**I**) Dotplot showing mean expression of marker genes and percentage of cells expressing them for each level 3 endothelial cell type. (**J**) Beeswarm plot of the log fold changes in Milo cell neighbourhoods in endothelial cells by CLD status. Milo neighbourhoods grouped into each level 3 endothelial cell type subcluster. Neighbourhoods with a significant (FDR <0.05) change in cellular abundance are coloured, with blue representing enrichment in healthy and red denoting expansion in CLD. Black lines denote mean logFC for each cell type.

Several groups have previously shown that SAMacs expand in CLD^14,15^ and other fibrotic diseases^15,43^. We performed differential abundance analysis on the MP atlas using the Milo package^44^, demonstrating that both SAMac1 and SAMac2 expanded in CLD livers when compared to healthy (Fig. 2C, D). We also used Welsh’s two sample t-tests as an alternative statistical approach to confirm the significant expansion of both SAMac1 and SAMac2 populations in CLD livers (Extended Data Fig. 4C). No differences were observed in MP composition between livers of male and female patients (Extended Data Fig. 4D). Interestingly, the expansion of SAMac2 was consistent across all aetiologies of CLD, whilst SAMac1 expansion was more notable in patients with biliary disorders PBC (primary biliary cholangitis) and PSC (primary sclerosing cholangitis) (Extended Data Fig. 4E). These data suggest that expansion of the SAMac2 population is a common pathological feature of CLD.

### Comprehensive annotation of other hepatic cell types identifies disease-associated populations

To identify other CLD-associated cell states, we replicated the level 3 analysis for each of the other major cell lineages defined in the atlas. A total of 7,301 mesenchymal cells were partitioned into 8 clusters (Fig. 2E), with distinct marker gene profiles (Fig. 2F; Supplementary Table 6). Specifically, we were able to resolve 4 distinct subpopulations of *MYH11*+ vascular smooth muscle cells (VSMCs), *LHX2*+ *NR1H4*+ quiescent and activated hepatic stellate cells (HSC), *THY1*+ *FBLN1*+ *IGF1*+ portal fibroblasts and *MSLN*+ mesothelial cells (Fig. 2F). Differential abundance analysis using Milo, demonstrated expansion of the portal fibroblast subpopulation in patients with CLD (Fig. 2G). This population, along with the activated HSCs, expressed the highest levels of fibrillar collagens (*COL1A1*, *COL1A2*, *COL3A1*) (Fig. 2F), suggesting they are the main scar-producing populations in the CLD liver.

Profound changes in the vasculature are also a characteristic feature of advanced CLD and regulates fibrosis^14,45,46^. 36,591 liver endothelial cells were analysed in our level 3 atlas, partitioning into 10 clusters (Fig. 2H). Identification of distinguishing marker genes (Fig. 2I, Supplementary Table 7) enabled annotation of liver sinusoidal endothelial cells (LSEC), central vein endothelia (CVEndo) enriched for *RSPO3*, *FBLN5*+ hepatic artery endothelia (HAEndo), *CCL21*+ *PDPN*+ lymphatic endothelia (Lymph Endo) as well as 3 additional subpopulations of vascular endothelial cells (Vasc endo 1-3). Additionally, we confirmed the presence of *PLVAP* and *ACKR1*+ scar-associated endothelial cells (SAEndo), as previously described^14^. Milo analysis demonstrated significant changes in endothelial cell composition in CLD, with expansion of arterial, lymphatic and SAEndo subpopulations, as well as a reduction in homeostatic sinusoidal and vascular subpopulations (Fig. 2J).

Focussing on lymphoid cells, we were able to resolve 19 subpopulations of T/NK cells (Extended Data Fig. 5A), which were manually annotated based on marker gene expression and prior literature (Extended Data Fig. 5B, Supplementary Table 8). Identified hepatic T cell subpopulations included naïve, memory, tissue resident memory (TRM) and regulatory (Treg) CD4+ T cells (Extended Data Fig. 5B). Similar heterogeneity was identified in the CD8+ T cell compartment, with distinct subpopulations of naïve, effector memory (TEM), resident memory and cytotoxic cells (Extended Data Fig. 5B). We also resolved subpopulations *GNLY*+ CD56^dim^ NK cells, *CD160*+ *EOMES*+ CD56^bright^ NK cells and 3 clusters of *SLC4A10*+ *IL23R*+ mucosal-associated invariant T (MAIT) cells (Extended Data Fig. 5B). Differential abundance analysis showed a striking change in T/NK cell composition in CLD, with loss of NK and cytotoxic CD8+ T cells and shift towards a CD4 rich environment with expansion of CD4+ naïve, CD4+ TRM and Tregs (Extended Data Fig. 5C).

Heterogeneity of the hepatic B cell compartment was also observed in the level 3 atlas (Extended Data Fig. 5D), with distinct subpopulations of *TCL1A*+ *IL4R*+ naïve B cells, *NR4A2*+ *TNFRSF13B*+ memory B cells and 2 clusters of *MZB1*+ *SDC1*+ plasma cells (Extended Data Fig. 5E, Supplementary Table 9). All naïve and memory B cell subpopulations were broadly expanded in CLD livers (Extended Data Fig. 5F). This expansion was most striking for *IL6*+ *MAST4*+ activated memory B cells (Extended Data Fig. 5E-F).

Finally, we utilised our level 3 atlas to annotate epithelial (hepatocyte and cholangiocyte) subpopulations and identify patterns of gene expression in hepatocytes indicative of metabolic zonation (Extended Data Fig. 6A-B, Supplementary table 10). Hepatocytes were mainly derived from the studies by Macparland and Andrews (Extended Data Fig. 6C), so robust conclusions about CLD-associated changes in hepatocyte composition or transcription cannot be drawn from this atlas. Cholangiocytes were captured from all the included studies (Extended Data Fig. 6C).

In summary, this atlas allows a broad understanding of changes in cellular composition of liver non-parenchymal cells in CLD.

### OLR1+ SAMac2 have a more pro-inflammatory and pro-fibrotic transcriptional profile

Having identified 2 distinct populations of *TREM2*+ *CD9*+ SAMac which expand in cirrhosis, we proceeded to define the transcriptional differences between SAMac1 and SAMac2. Differential expression analysis demonstrated transcriptional heterogeneity between these populations, with 903 genes significantly enriched in SAMac1 and 1176 genes in SAMac2 (Fig. 3A, Supplementary table 11). Specifically, SAMac1 expressed higher levels of the previously described SAMac marker *GPNMB*^15^ as well as several anti-inflammatory anti-fibrotic factors such as *MMP9*, *IL10, GDF15, CD36* and *PPARG* (Fig. 3B). In contrast, SAMac2 demonstrated higher expression of pro-inflammatory and pro-fibrotic mediators (*IL1B*, *TNF*, *IL18*, *NLRP3*, *AREG*, *PDGFB*) as well as scavenger receptors including *CLEC5A* and *OLR1* (Fig. 3B). Using OLR1 as a marker of SAMac2, we employed multiplex immunofluorescent staining on human CLD tissue to confirm that OLR1+ CD68+ cells were spatially located in areas of fibrosis and represented a subpopulation of CD68+ SAMacs (Fig. 3C and Extended Data Fig. 7A). OLR1+ CD68+ SAMac2 (Fig. 3D; red arrows) were distinct from GPNMB+ CD68+ SAMac1 (Fig. 3D; yellow arrows), further highlighting SAMac heterogeneity in the human liver fibrotic niche and demonstrating the potential utility of OLR1 and GPNMB as markers of SAMac subpopulations.

**Figure 3:**
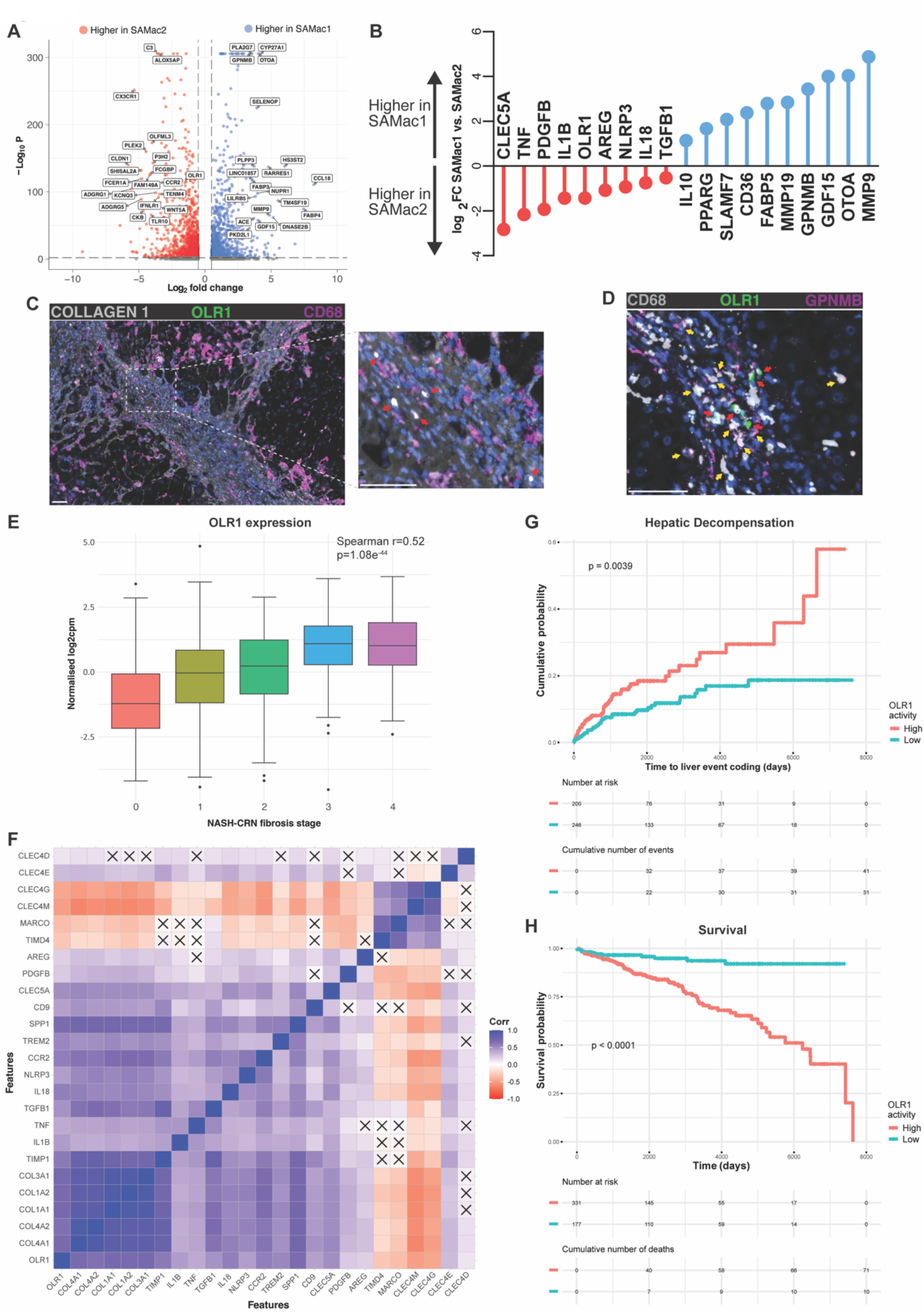
High OLR1 expression distinguishes pro-inflammatory SAMacs and is associated with CLD progression. (**A**) Volcano plot of differential gene expression between SAMac1 and SAMac2 in the level 3 MP atlas. Significant genes (Log_2_FC>0.5 and adjusted *p*-value<0.01) coloured in blue if enriched in SAMAc1 and red if enriched in SAMac2. Selected genes are labelled. (**B**) Log_2_FC plot of selected significant genes from A. Circles in red are genes enriched in SAMac2 and circles in blue enriched in SAMac1. (**C**) Representative multiplex immunofluorescence image of human cirrhotic liver at low magnification and high magnification of the fibrotic niche. Markers are Collagen 1 (grey), CD68 (magenta), OLR1 (green) and DAPI (blue). Scale bar=50μm. Red arrows indicate OLR1+ CD68+ SAMacs. (**D**) Representative multiplex immunofluorescence image of the fibrotic niche of human cirrhotic liver. Markers are CD68 (grey), GPNMB (magenta), OLR1 (green) and DAPI (blue). Scale bar=50μm. Red arrows indicate OLR1+ CD68+ SAMacs. Yellow arrows indicate GPNMB+ CD68+ SAMacs. (**E**) Box and whiskers plot of OLR1 gene expression (shown as normalised log_2_ counts per million (CPM)) according to histological fibrosis stage (NASH-CRN) in 632 cases (n=172 (F0), 134 (F1), 96 (F2), 111 (F3), 119 (F4)) from the SteatoSITE data set. All boxplots show median (centre line), first and third quartiles (lower and upper box limits), 1.5× interquartile range (whiskers) and outliers (dots). Spearman correlation of normalised OLR1 cpm vs ordinal NASH-CRN F-stage of cases was calculated. (**F**) Correlation matrix of normalized gene expression counts for stated genes in the complete n = 663 set of cases (including controls) with RNA-seq from the SteatoSITE data set. Statistically significant correlations (*p* < 0.05) shown as shaded boxes and nonsignificant correlations marked with an X. Boxes coloured by Spearman correlation coefficients (blue=positive and red=negative correlation). (**G**) Biopsy cases from the SteatoSITE data set (n = 446) with available RNA-seq and no hepatic decompensation– related coding event before the time of the biopsy were used for time-to-event analysis. Normalized gene expression counts for OLR1 were used to divide cases into low and high risk of hepatic decompensation (Kaplan-Meier estimator curves with log-rank test *p-*value shown). (**H**) Biopsy cases from the SteatoSITE dataset (n = 508) with available RNA-seq were used for time-to-all cause mortality analysis. Normalized gene expression counts for OLR1 was used to divide cases into low and high risk of death (Kaplan–Meier estimator curves with log-rank test *p*-value shown).

### High OLR1 expression is associated with liver disease progression and adverse clinical outcomes

OLR1 (LOX-1), a class E scavenger receptor, has been suggested as a therapeutic target for atherosclerosis^30^ with candidate compounds currently being evaluated in clinical trials in patients with cardiovascular disease^33^. *OLR1* expression, alongside *CLEC5A*, was specific to MPs in the human liver, highly enriched in SAMacs and not expressed significantly by other hepatic cell lineages (Extended Data Fig. 7B-C), suggesting it may offer a route to selective modulation of SAMacs without off target effects on other cells. Indeed, *OLR1* and *CLEC5A* were more specific than the majority of previously reported SAMac markers (*CD9*, *CD63*, *FABP5*, *SPP1*, *GPNMB*), with only *TREM2* showing similar levels of specificity (Extended Data Fig. 7B-C).

To determine whether OLR1 could represent a therapeutic target in CLD across the disease severity spectrum, we analysed expression in liver bulk RNA-seq data from MASLD liver samples in the SteatoSITE cohort^2^. This showed a significant correlation between *OLR1* expression and histological fibrosis stage in patients with MASLD (Fig. 3E). Furthermore, hepatic OLR1 expression demonstrated a significant positive correlation with other fibrogenic and inflammatory genes, and a negative correlation with genes expressed by homeostatic non-parenchymal cell populations such as KCs (*TIMD4*, *MARCO*) or LSECs (*CLEC4M*, *CLEC4G*) (Fig. 3F). The SteatoSITE cohort also includes linked clinical outcome data^2^. Strikingly, high hepatic *OLR1* expression was associated with increased rates of liver disease progression (Fig. 3G) and higher overall mortality (Fig. 3H) in patients with MASLD. Overall, these data indicate that OLR1+ SAMacs may promote hepatic inflammation and fibrogenesis across the full spectrum of CLD severity.

### OLR1+ scar-associated monocyte-derived macrophages are conserved across species

To assess whether a similar pro-inflammatory OLR1+ SAMac subpopulation is conserved across species, we employed the well characterised chronic carbon tetrachloride (CCl_4_) mouse model of liver fibrosis progression and regression^18^. We performed scRNAseq on blood and liver CD45^+^ immune cells from uninjured (healthy) control mice, mice with liver fibrosis (24 and 96hrs) and mice following fibrosis regression (4 weeks) (Fig. 4A). Following QC and lineage annotation (Extended Data Fig. 8A-B), a total of 18,177 MPs from blood and liver were annotated for downstream analysis. Clustering of mouse MPs revealed 17 clusters of cells derived from liver and blood of mice across the different conditions (Fig. 4B). Manual annotation was performed using previously described marker genes (Fig. 4C, Supplementary table 12), identifying the expected subpopulations of monocytes, macrophages and DCs. We also identified previously unreported mouse MP subpopulations, including a specific liver macrophage cluster which expressed high levels of *Acod1* and *Cxcl10* (annotated as MDM Acod1+) (Fig. 4B-C).

**Figure 4:**
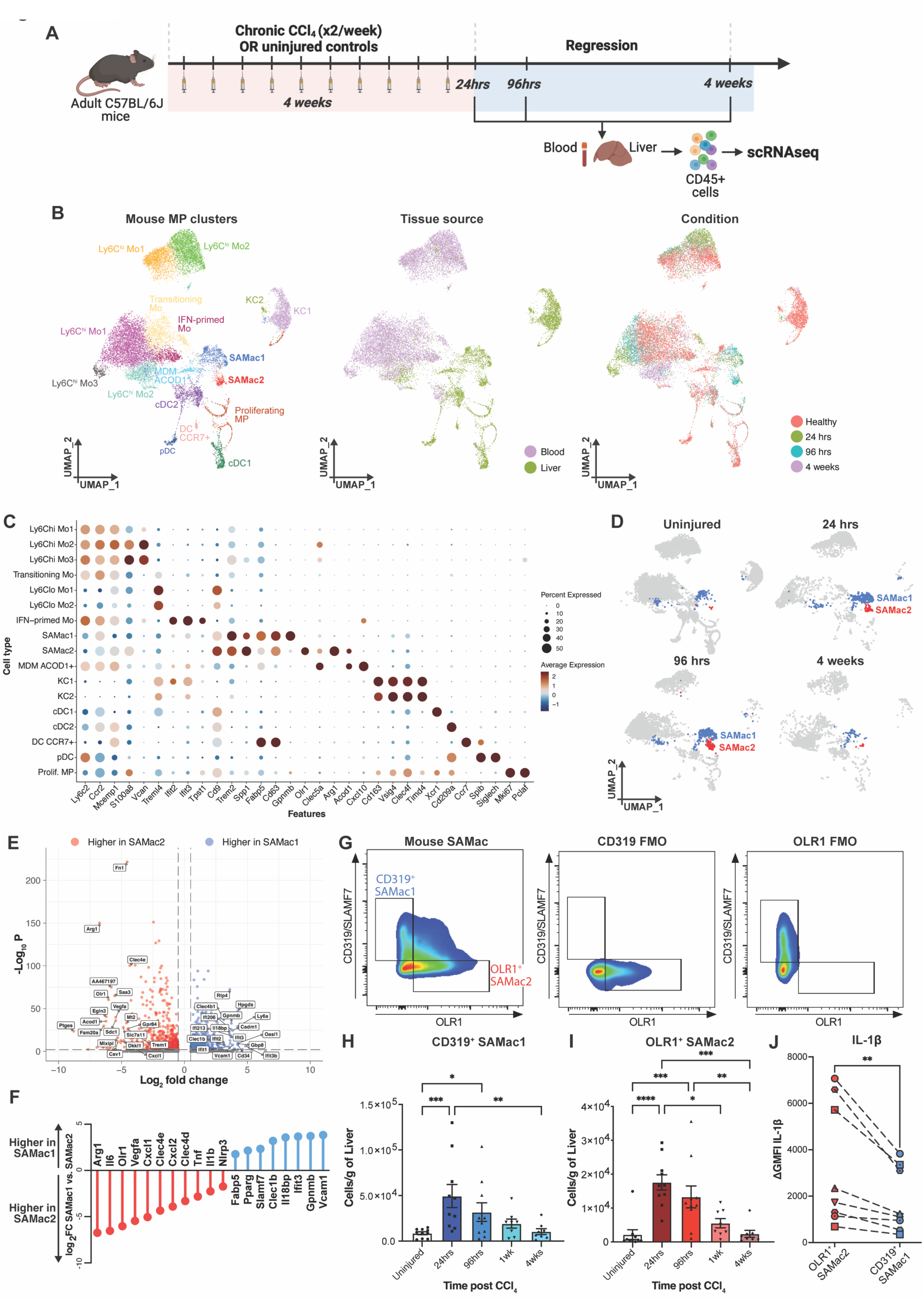
Pro-inflammatory OLR1+ SAMacs accumulate in mouse CLD. (**A**) Overview of experimental workflow for mouse scRNAseq data generation. Created in BioRender. Colella, F. (2025) https://BioRender.com/su1qjok. (**B**) UMAPs of mouse MP scRNAseq data (18,177 cells). Coloured and labelled by MP cell types annotation (right). Coloured by tissue source blood vs liver (middle). Coloured by condition/CCl4 model timepoint (right). (**C**) Dotplot showing mean expression of marker genes and percentage of cells expressing them for each mouse MP cell type. (**D**) UMAPs of mouse MP scRNAseq data split by stated condition. SAMac1 labelled in blue and SAMac2 labelled in red. Other MP cell types labelled in grey. (**E**) Volcano plot of differential gene expression between SAMac1 and SAMac2 in the mouse MP atlas. Significant genes (Log_2_FC> 0.5 and adjusted *p*-value<0.01) coloured in blue if enriched in SAMac1 and red if enriched in SAMac2. Selected genes are labelled. (**F**) Log_2_FC plot of selected significant genes from E. Circles in red are genes enriched in SAMac2 and circles in blue enriched in SAMac1. (**G**) Representative flow cytometry plots showing gating strategy for identification of OLR1+ SAMac2 and CD319+ SAMac1 in mouse liver. Samples pregated on viable mouse liver SAMacs (Extended Data Fig. 9A). Full stained, CD319 fluorescence minus one (FMO) and OLR1 FMO controls shown. (**H**) Bar plot of CD319+ SAMac numbers (quantitated using flow cytometry) in livers of uninjured (control) mice and in mice at stated timepoint following 4 weeks of chronic CCl_4_ administration. (**I**) Bar plot of OLR1+ SAMac numbers (quantitated using flow cytometry) in livers of uninjured (control) mice and in mice at stated timepoint following 4 weeks of chronic CCl_4_ administration. For H and I, bars indicate mean ± S.E.M. with individual dots representing independent mice. Kruskal-Wallis test with Dunn’s multiple comparison performed; \**p*<0.05, \*\**p*<0.01, \*\*\**p*<0.001, \*\*\*\**p*<0.0001. (**J**) IL-1β production in OLR1+ SAMac2 and CD319+ SAMac1 isolated from chronic CCl_4_ treated mouse livers was assessed by intracellular flow cytometry. Shapes represent independent mice, with SAMac2 coloured in red and SAMac1 coloured blue for each mouse. To measure active IL-1β production, Δ geometric mean fluorescence intensity (ΔGMFI) of IL-1β was calculated for SAMac1 and SAMac2 for each sample, by subtracting the IL-1β GMFI of control cells (incubated with momensin at 4°C for 3 hrs) from the IL-1β GMFI following incubation with momensin at 37°C for 3 hours. Wilcoxon matched-pairs signed rank test performed; \*\**p*<0.001.

Focussing on the SAMacs, similar to human CLD, we identified 2 subpopulations of *Trem2*+ *Cd9*+ *Cd63*+ *Spp1*+ SAMacs, termed SAMac1 and SAMac2 (Fig. 4B-C). Both mouse SAMac subpopulations were derived from the liver samples (Fig. 4B) and expanded at fibrotic timepoints (24 and 96 hrs) but had returned to baseline levels 4 weeks following cessation of injury (Fig. 4D). As with human SAMacs, differential expression analysis between mouse SAMac1 and SAMac2 identified significant transcriptional heterogeneity (Fig. 4E, Supplementary Table 13), with SAMac1 being enriched for *Gpnmb*, *Slamf7* and other anti-inflammatory mediators (e.g. *Il18bp*, *Pparg*), whilst mouse SAMac2 showed elevated expression of *Olr1* and pro-inflammatory factors including inflammasome components *Il1b* and *Nlrp3* (Fig. 4F). Hence, our finding of SAMac heterogeneity is conserved between human and mouse CLD.

A publicly accessible browser for mouse MP data is available at: https://shiny.igc.ed.ac.uk/mouseMPshinyApp/.

We proceeded to study the kinetics of mouse SAMac1 and 2 in the CCl_4_ liver fibrosis model. A flow cytometry panel was developed to enable the distinction of CD319/Slamf7+ SAMac1 and Olr1+ SAMac2 (Extended Data Fig. 9A, Fig. 4G). Both Olr1+ SAMacs and CD319+ SAMacs expanded significantly during active liver injury but had returned to baseline levels 4 weeks after the cessation of injury (Fig. 4H-I). To further validate our transcriptional findings that Olr1+ SAMacs have a more pro-inflammatory phenotype, we assessed *ex vivo* pro-inflammatory cytokine production via intracellular flow cytometry on macrophages isolated from fibrotic mouse liver. In keeping with our predictions from the transcriptional data, Olr1+ SAMacs demonstrated increased production of IL-1β when compared to CD319+ SAMacs (Fig. 4J).

Previous pseudotemporal trajectory analysis and fate mapping have demonstrated that SAMacs overall derive from circulating monocytes rather than resident KCs^14,47,48^. To confirm the origin of both SAMac subpopulations we generated busulfan conditioned bone-marrow chimeric mice^49^, with CD45.1+ expression used to track bone-marrow derived donor cells into the liver in the context of CLD (Extended Data Fig. 9B). A flow cytometry gating strategy was developed to study the degree of donor cell contribution to liver myeloid subpopulations including Ly-6C^hi^ monocytes, Timd4^+^ Vsig4^+^ resident KCs, Timd4^-^Vsig4^+^ monocyte-derived KCs (MoKCs) and the 2 subpopulations of SAMacs (Extended Data Fig. 9C). Notably, using this system there was minimal replenishment of resident KCs under homeostatic conditions (Extended Data Fig. 9D). Following chronic injury with CCl_4_, Timd4^+^ resident KCs remained mainly of host origin, suggesting there is little monocyte contribution to this population (Extended Data Fig. 9E). In contrast, SAMacs and MoKCs were entirely derived from donor bone marrow, with no detectable difference in chimerism between the 2 SAMac subpopulations (Extended Data Fig. 9E). Hence, both subpopulations of SAMacs are derived from the recruitment and differentiation of circulating monocytes and not from resident KCs.

### OLR1+ scar-associated macrophages are enriched for pro-inflammatory transcription factor motifs

To assess the transcriptional regulation of SAMac subpopulations in CLD, we performed single-cell ATAC-seq (scATACseq) on blood and liver CD45+ immune cells from uninjured mice and mice treated with chronic CCl_4_ during fibrogenesis (24hrs) and following regression (4 weeks) (Fig. 5A). Following initial QC and clustering of blood and liver scATACseq data (Extended Data Fig. 10A), cell lineages were computationally annotated via a label transfer from scRNAseq data (Extended Data Fig. 10B-C) and MPs subsetted for downstream analysis. Reclustering MPs revealed 15 scATACseq clusters from blood and liver (Fig. 5B), across the different conditions (Extended Data Fig. 10C). ScATACseq MP clusters were then computationally annotated by integrating with the mouse MP scRNAseq dataset and transferring labels from RNA to ATAC clusters (Fig. 5C). This generated an annotated mouse MP scATACseq dataset (Fig. 5D). Notably, similar to the human and mouse scRNAseq data, we were able to distinguish SAMac1 and SAMac2 clusters based on scATACseq clustering, indicating that these populations have distinct chromatin accessibility profiles (Fig. 5D).

**Figure 5:**
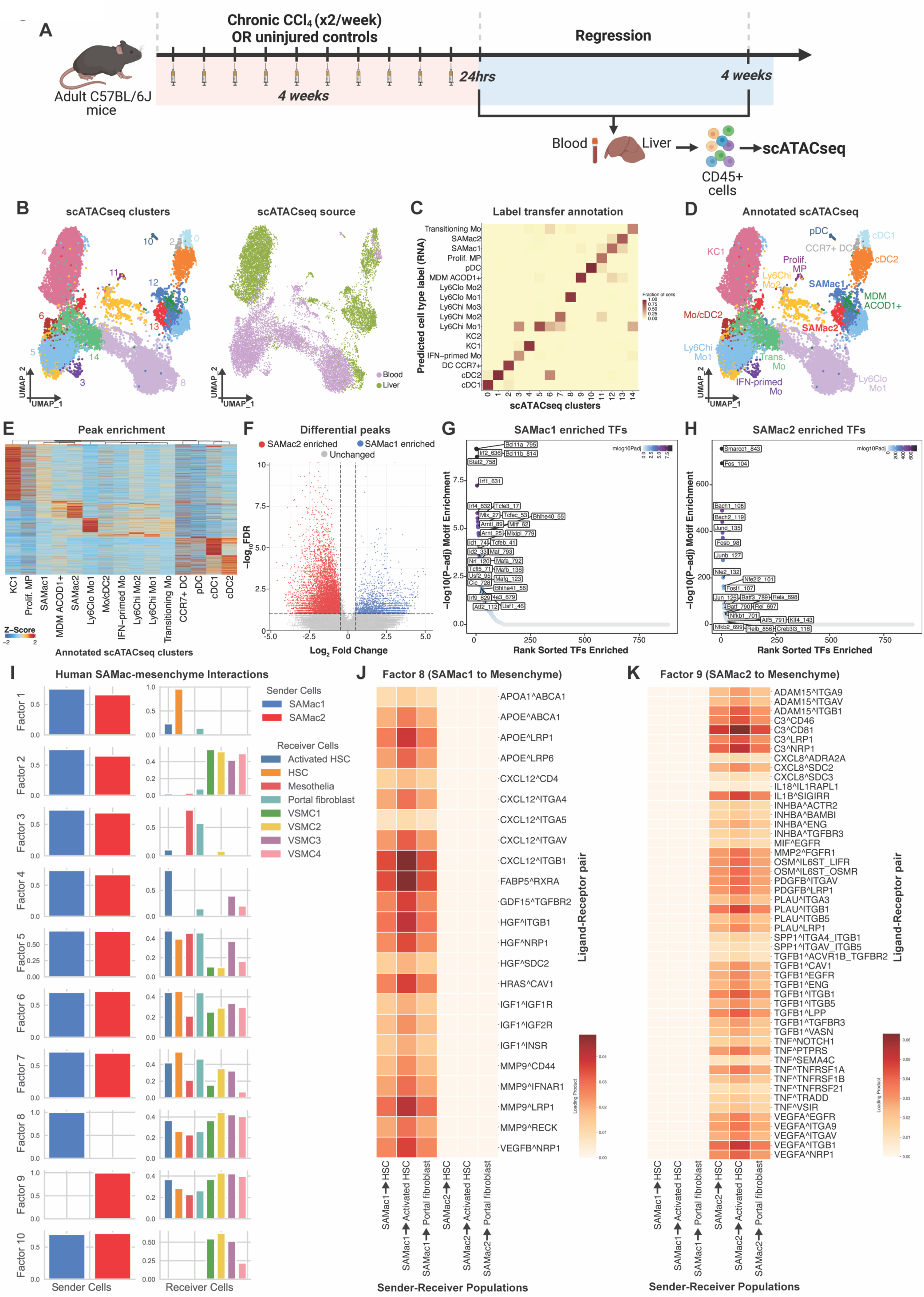
OLR1+ SAMacs are enriched for pro-inflammatory transcription factor motifs and cell-cell interactions with mesenchymal cells. (**A**) Overview of experimental workflow for mouse scATACseq data generation. Created in BioRender. Colella, F. (2025) https://BioRender.com/xzgt2un. (**B**) UMAPs of mouse MP scATACseq data (14, 613 cells). Coloured and labelled by scATACseq clusters (left) and by tissue source blood vs liver (right). (**C**) Heatmap of label transfer annotation analysis between mouse MP scATACseq clusters and annotated mouse MP scRNAseq clusters (RNA). Colour scale indicates prediction score of annotation of each scATACseq cluster. The highest prediction scores were used to annotate scATACseq clusters. (**D**) UMAP of mouse MP scATACseq data coloured and labelled by cell type annotation. (**E**) Heatmap representation of marker chromatin accessibility peak enrichment in each annotated MP cell type from the scATACseq peak set. Each row represents a peak, each column represents an annotated cluster in the scATACseq dataset. Colour scale indicates Z-score. (**F**) Volcano plot of differential peak enrichment between SAMac1 and SAMac2 in the mouse scATACseq MP data. Significant peaks (Log_2_FC> 0.5 and FDR<0.1) coloured in blue if enriched in SAMac1 and red if enriched in SAMac2. (**G**) Rank significance plot of transcription factor (TF) motifs enriched in SAMac1 vs SAMac2 from scATACseq data. Selected TF motifs labelled. TF motifs coloured by significance. (**H**) Rank significance plot of transcription factor (TF) motifs enriched in SAMac2 vs SAMac1 from scATACseq data. Selected TF motifs labelled. TF motifs coloured by significance. (**I**) Ligand-receptor interaction analysis in human level 3 atlas, between SAMac1 and SAMac2 (as senders) and mesenchymal cell clusters as receivers. 10 factors were identified representing a different cell-cell communication programs. Loadings (y-axis) for different sender and receiver cell types are shown for each factor. (**J**) Heatmap of selected ligand-receptor pairs in factor 8. Rows represent ligand-receptor pairs; columns represent specific cell type interactions; colour scale represents the loading product (degree of enrichment). (**K**) Heatmap of selected ligand-receptor pairs in factor 9. Rows represent ligand-receptor pairs; columns represent specific cell type interactions; colour scale represents the loading product (degree of enrichment).

To study candidate transcription factors (TFs) which regulate SAMac responses, we proceeded to interrogate the open chromatin profiles from the annotated MP scATACseq data. Mouse MP subpopulations demonstrated distinct accessible peak profiles (Fig. 5E), with significant differences observed when comparing peak accessibility between SAMac1 and SAMac2 (Fig. 5F). TF motif analysis on SAMac1 enriched peaks revealed enrichment for *Bcl11A* and *Bcl11B* motifs, as well as several interferon responsive TFs including *Irf1*, *Irf2*, *Irf4* and *Stat2* (Fig. 5G, Supplementary table 14). In contrast, Olr1+ SAMac2 showed enrichment for several pro-inflammatory TF motifs including *Fos*, *Fosb*, *Jun, NfkB* and family members *Rel* and *Rela* (Fig. 5H, Supplementary table 14). Interestingly, SAMac2 was also enriched for several transcriptional repressors such as *Nfe2*, *Nfe2l2* and *Bach2*, suggesting a balance between activation and regulatory signals (Fig. 5H). Hence, in addition to a pro-inflammatory transcriptome, Olr1+ SAMac2 have an open chromatin profile which is primed for inflammatory responses in the injured liver.

### OLR1+ scar-associated macrophages are enriched for pro-inflammatory interactions with scar-producing mesenchyme

Having determined that OLR1+ SAMac2 has a more pro-inflammatory pro-fibrogenic transcriptional profile than GPNMB+ SAMac1 in both human and mouse liver fibrosis, we then sought to assess whether they have distinct interactions with mesenchymal cells which would favour activation and ECM deposition. Using the LIANA+ package^50^, we performed cellular interactome analysis between the 2 SAMac subpopulations (as senders) and mesenchymal cell clusters (as receivers) from our level 3 atlas. Focussing on HSCs, activated HSCs and portal fibroblasts as the main mesenchymal cells which express ECM and regulate liver fibrosis, we identified 3 groups (factors) of ligand-receptor pairs where ligands were expressed by both SAMac1 and SAMac2 (Fig. 5I, Supplementary table 15). Factor 1 incorporated ligand-receptor interactions predominantly between SAMac1/2 and HSCs and included galectins and TIMPs (Fig. 5I, Extended Data Fig. 10E). Factor 4 interactions were mainly between SAMac1/2 and activated HSCs and included calmodulins and osteopontin (*SPP1*) as ligands (Fig. 5I, Extended Data Fig. 10F). Factor 3 interactions were enriched between both SAMac1 and SAMac2 as the senders and portal fibroblasts as the receiver population and included ligands proganulin (*GRN*) and macrophage migration inhibitory factor (*MIF*) (Fig. 5I, Extended Data Fig. 10G).

Our interactome analysis also identified 2 factors which were specific to SAMac1 and SAMac2 as senders, factor 8 and factor 9 respectively (Fig. 5I, Supplementary table 15). Factor 8 (SAMac1) ligands included several reported anti-fibrotic factors including *HGF*^51^, *IGF1*^52^, *MMP9*^53^ and *GDF15*^54^ (Fig. 5J). In contrast, factor 9 (SAMac2) ligands included *TNF*^55^, *TGFB1*, *IL1B*^25,27,28^, *IL18*, *C3*^53^ and *PDGFB* (Fig. 5K), mediators which have been associated with promoting mesenchymal cell activation and fibrogenesis in the liver and/or other disease contexts. These data further highlight that the transcriptional heterogeneity in SAMacs has potential consequences for signalling to fibrogenic mesenchymal cells, with OLR1+ SAMac2 predicted to promote and propagate activation of scar-producing activated HSCs and portal fibroblasts.

### OLR1+ macrophages promote fibrosis in multilineage human liver spheroids

To determine whether OLR1+ monocyte-derived macrophages have a functional role in liver fibrosis, we developed a human multilineage spheroid culture system, containing embryonic stem (ES) cell-derived hepatocytes, hepatic stellate cells and endothelial cells as previously described^56^, coupled with the addition of primary human monocytes from patients with CLD (Fig. 6A). The addition of monocytes, followed by 7 days of monocyte differentiation and spheroid maturation *in vitro*, resulted in engraftment of IBA1+ MDMs embedded within the spheroids (Fig. 6B). The addition of patient monocytes increased spheroid expression of SAMac2 markers *OLR1*, *SPP1* and *CD68,* and promoted spontaneous fibrogenic activity with higher levels of *COL1A1* following 1 week of maturation (Fig. 6C). No alteration was observed in hepatocyte gene *TTR*. To confirm that this spheroid system could be utilised to test the antifibrotic effects of drug therapy, the TGFβ signalling inhibitor A83-01 or vehicle control were administered to spheroids during the treatment phase, between day 8 and day 15 (Fig. 6A). TGFβ inhibition attenuated fibrogenic gene expression (*COL1A1* and *COL3A1*) in macrophage containing spheroids (Extended Data Fig. 11A). We then proceeded to test the functional impact of OLR1+ macrophage blockade, via administration of the CSF1R inhibitor Pexidartinib to spheroids during the treatment phase. CSF1R blockade significantly reduced *OLR1*, *IL1B*, *SPP1* and *CD68* and attenuated fibrillar collagen (*COL1A1* and *COL3A1)* gene expression (Fig. 6D), confirming that human OLR1^+^ MDMs have a pro-fibrotic function.

**Figure 6:**
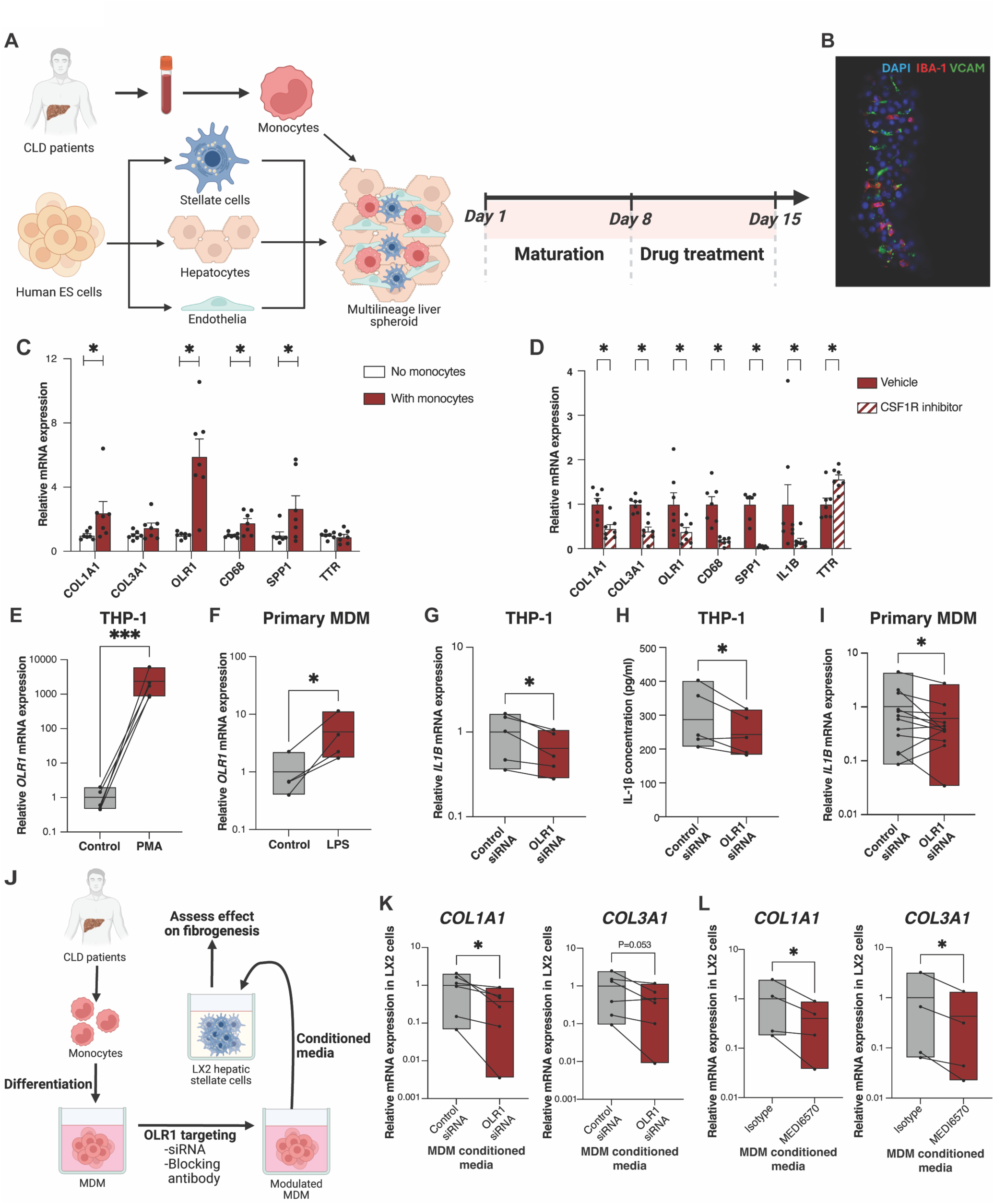
Targeting OLR1+ monocyte-derived macrophages attenuates inflammatory and fibrogenic activity. (**A**) Overview of experimental workflow for generating embryonic stem (ES) cell derived multilineage spheroids containing primary human monocyte-derived macrophages from CLD patients. Following aggregation and addition of monocytes on Day 1, spheroids mature for 7 days, followed by an optional drug treatment phase between day 8 (D8) and 15 (D15). Created in BioRender. Colella, F. (2025) https://BioRender.com/sp1zyj3. (**B**) Representative multiplex immunofluorescence of a D8 human liver spheroid following monocyte addition. Markers are IBA1 (red), VCAM1 (green) and DAPI (blue). (**C**) qPCR of D8 spheroids comparing aggregation with and without addition of patient monocytes. Dots represent biological replicates with monocytes from different CLD patients. Expression of stated gene in each spheroid culture relative to mean expression of control (no monocyte) group is shown. Bars indicate mean ± S.E.M. for each group. Wilcoxon matched-pairs signed rank test performed between groups for each gene; \**p*<0.05. (**D**) qPCR of D15 spheroids comparing 7 days treatment with CSF1R inhibitor (Pexidartinib) or vehicle control. Dots represent biological replicates with monocytes from different CLD patients. Expression of stated gene in each spheroid culture relative to mean expression of control (vehicle) group is shown. Bars indicate mean ± S.E.M. for each group. Wilcoxon matched-pairs signed rank test performed between groups for each gene; \**p*<0.05. (**E**) Relative *OLR1* mRNA expression of THP1 cells following treatment with PMA or vehicle control. Bars are min to max expression with line at mean. Dots connected by lines represent data from 5 independent experiments. Ratio paired t-test, \*\*\**p*<0.001. (**F**) Relative *OLR1* mRNA expression of primary MDM following treatment with LPS or vehicle control. Bars are min to max expression with line at mean. Dots connected by lines represent data from 4 CLD patients. Ratio paired t-test, \**p*<0.05. (**G**) Relative *IL1B* mRNA expression in differentiated THP1 cells following treatment with OLR1 targeting or control siRNA. Bars are min to max expression with line at mean. Dots connected by lines represent data from 5 independent experiments. Ratio paired t-test, \**p*<0.05. (**H**) IL-1β concentration measured by ELISA in the supernatant of differentiated THP1 cells following treatment with OLR1 targeting or control siRNA. Bars are min to max expression with line at mean. Dots connected by lines represent data from 5 independent experiments. Ratio paired t-test, \**p*<0.05. (**I**) Relative *IL1B* mRNA expression of primary MDM following treatment OLR1 targeting or control siRNA. Bars are min to max expression with line at mean. Dots connected by lines represent data from 12 different CLD patients. Wilcoxon matched-pairs signed rank test, \**p*<0.05. (**J**) Overview of experimental workflow for assessing effect of MDM OLR1 modulation on HSC fibrogenic activity. Created in BioRender. Colella, F. (2025) https://BioRender.com/l42u16a. (**K**) Relative *COL1A1* and *COL3A1* mRNA expression of LX2 hepatic stellate cells following incubation with conditioned media from primary MDMs pre-treated with OLR1 targeting or control siRNA. Bars are min to max expression with line at mean. Dots connected by lines represent data from MDMs of 6 different CLD patients. Ratio paired t-test, \**p*<0.05. (**L**) Relative *COL1A1* and *COL3A1* mRNA expression of LX2 hepatic stellate cells following incubation with conditioned media from primary MDMs pre-treated with OLR1 blocking antibody (MEDI6570) or isotype control antibody. Bars are min to max expression with line at mean. Dots connected by lines represent data from MDMs of 4 different CLD patients. Ratio paired t-test, \**p*<0.05.

### Targeting OLR1 reduces inflammatory and fibrogenic activity in human macrophages

Having demonstrated that hepatic OLR1+ SAMacs have pro-inflammatory and pro-fibrogenic phenotype, we then sought to determine whether OLR1 itself represents an anti-inflammatory therapeutic target for liver fibrosis. OLR1 expression was upregulated in THP-1 MDMs in response to PMA (phorbol 12-myristate 13-acetate) treatment (Fig. 6E), and in primary human MDMs in response to LPS treatment (Fig. 6F), indicating that macrophage OLR1 expression is induced by pro-inflammatory stimuli. Knockdown of OLR1 using siRNA in THP-1 MDMs resulted in less *OLR1* gene expression (Extended Data Fig. 11B), reduced *IL1B* gene expression (Fig. 6G) and attenuated IL-1β protein secretion (Fig. 6H). Similarly, treatment of primary human MDMs from CLD patients with OLR1 siRNA reduced *OLR1* gene expression (Extended Data Fig. 11C) and significantly decreased *IL1B* gene expression (Fig. 6I).

We proceeded to test the impact of human macrophage OLR1 modulation on fibrogenic activity of HSCs (Fig. 6J). Conditioned media from primary MDMs with OLR1 knockdown was less able to induce fibrogenic activity in LX2 HSCs than control conditioned media (Fig. 6K). Furthermore, treatment of primary human MDMs with MEDI6570, a clinically used OLR1 blocking antibody^33^, attenuated fibrillar collagen expression in LX2 cells following conditioned media transfer (Fig. 6L). No effect was observed on LX2 survival with OLR1 knockdown or antibody blockade of MDMs (Extended Data Fig. 11D-E). These data highlight that targeting OLR1 in MDMs can inhibit inflammatory mediator production and reduce signalling to ECM-producing myofibroblasts, attenuating fibrogenic activity. Hence, OLR1 inhibition is a potential novel therapeutic approach to selectively modulate scar-associated macrophage phenotype and inhibit liver fibrogenesis.

## Discussion

Our annotated human CLD cellular atlas, comprising over 600,000 cells from 77 patients, has revealed previously unrecognised liver cell heterogeneity and started to facilitate the identification of refined therapeutic targets for CLD. Going forwards, we anticipate this will be a valuable resource for the community to dissect the pathobiology of liver fibrosis and understand how this might be modulated. We focussed on the immune compartment, identifying a previously undescribed subpopulation of SAMacs which expands across different etiologies of human CLD, are conserved across species, and have a pro-inflammatory phenotype. OLR1 is a specific marker of these SAMacs and regulates inflammatory and fibrogenic activity in this population. Hence, targeting OLR1 represents a novel therapeutic approach for liver inflammation and fibrosis, while hepatic OLR1 levels could serve as a prognostic biomarker to improve risk stratification of patients with CLD.

SAMacs have been shown to expand in fibrotic human liver in several previous studies. However, data on their functional role in fibrosis is conflicting. Some studies have demonstrated a pro-fibrotic role for SAMacs via interactions with HSCs that promote activation and ECM deposition^14,15^. In contrast, other studies have suggested that TREM2+ SAMacs have an anti-inflammatory and anti-fibrotic function in CLD, promoting scar resolution and the restitution of normal tissue architecture^20–23^. It remains unclear whether these apparently opposing functions could be mediated by distinct subpopulations of SAMacs. Our human scRNAseq atlas, containing vastly more myeloid cells derived from more patients than previous analyses, has enabled us to identify human SAMac heterogeneity which could explain this functional dichotomy. In particular, the OLR1+ SAMac2 population displays a more pro-inflammatory and pro-fibrogenic phenotype, suggesting this may be the optimal target population to inhibit the pathogenic features of SAMacs in CLD.

To selectively modulate the pathogenic features of SAMacs without perturbing their pro-resolution functions, the ideal therapeutic target would be enriched specifically in the pro-inflammatory subpopulation and expressed on the cell surface to improve druggability. Having identified SAMac2 in our scRNAseq atlas, we screened this cell population for candidate targets which fulfil these therapeutic criteria. From our analyses, OLR1 (LOX-1) emerged as a potential target, reinforced by the strong association between elevated liver OLR1 expression and adverse clinical outcomes in the SteatoSITE cohort, and functional data demonstrating that targeting OLR1 on human MDMs reduces pro-inflammatory and pro-fibrogenic properties. OLR1 has already been explored as a therapeutic target in cardiovascular disease, with the specific OLR1 blocking monoclonal antibody MEDI6570 having been administered to patients in phase 1^33^ and phase 2 clinical trials (clinicaltrials.gov NCT04610892), with no safety concerns reported. This opens up the possibility of a drug repurposing study using MEDI6570 in CLD patients, potentially accelerating therapeutic development. The discovery of small molecule OLR1 inhibitors with anti-inflammatory effects should also be pursued, ideally with proven cross-reactivity in mice or other species, to facilitate *in vivo* validation of functional effects of OLR1 blockade on CLD pathogenesis, something which we have been unable to do using existing molecules. In addition, other candidate targets specifically expressed by pro-inflammatory SAMac2 are also worth exploring, for example CLEC5A which has been reported to mediate pro-inflammatory activity in myeloid cells^57^. Our identification and characterisation of human SAMac2 should therefore herald further studies aimed at identifying the optimal mechanistic approach to modulate this population.

A key feature of OLR1+ SAMac2 in both human and mouse CLD is elevated expression of inflammasome components including the pro-inflammatory cytokine IL-1β. Macrophage-derived IL-1β is known to promote mesenchymal cell activation and fibrogenesis in the liver^25^ and other human tissues such as heart^27,28^. However, while it has previously been unclear how to selectively attenuate macrophage IL-1β production in the fibrotic niche, our finding that targeting OLR1 on MDMs can reduce IL-1β production and fibrogenic activity suggests that OLR1 may potentially have a broader role as an anti-inflammatory target in fibrotic diseases in other tissues such as heart, lung and kidney which also show SAMac accumulation^29^. An integrated scRNAseq atlas and the application of high-resolution spatial transcriptomics across fibrotic human organs would allow a more detailed assessment of which specific diseases demonstrate OLR1 expression on IL-1β+ SAMacs. It will be important to include samples from different disease stages and etiologies in such analyses, to elucidate when in the disease trajectory targeting of pro-inflammatory SAMac will be most efficacious.

OLR1 has also been proposed as a prognostic biomarker for various types of cancer^32^ and for cardiovascular risk^30^. Our data suggest that hepatic OLR1 levels may also be a prognostic biomarker for CLD. One previous study suggested that elevated soluble OLR1 in the circulation could differentiate patients with MASLD from healthy controls, although the degree of fibrosis or macrophage accumulation in these patients was unclear^58^. While we have been unable to demonstrate differences in soluble OLR1 between healthy and advanced CLD patient serum using a commercial assay (data not shown), the potential for OLR1 as a prognostic biomarker should be investigated in larger patient cohorts using optimised immunoassays. The prognostic value of quantifying OLR1 (and/or other SAMac2 markers) in liver biopsy tissue via immunohistochemistry should also be assessed in patients with different causes of CLD. Ultimately, the accumulation of OLR1+ SAMacs may define subgroups of CLD patients who are more likely to respond to specific anti-inflammatory therapies targeting SAMac2.

Our findings of SAMac heterogeneity pose new questions about SAMac biology in the liver. While definitive fate mapping is not possible in humans, we have shown in mice that both SAMac1 and SAMac2 derive predominantly from circulating monocytes and not from resident KCs. Macrophages are plastic cells, capable of switching phenotype according to microenvironmental cues. It is unknown whether MDMs in the liver can switch between SAMac2 and SAMac1 phenotypes and which signals might regulate this process. Further *in vivo* experimental work using a time-controlled monocyte fate mapping system is needed to dissect the complexity of SAMac kinetics. We have highlighted OLR1 as a driver of inflammasome activation, IL-1β production and pro-fibrogenic activity of SAMac2. However, the ligands which activate OLR1 signalling in the CLD liver are yet to be defined. Oxidised LDL (oxLDL), the best described OLR1 ligand, accumulates in the CLD liver and could potentially drive SAMac2 activation and pro-inflammatory mediator production^59^. However, OLR1 is a pleiotropic scavenger receptor with numerous candidate ligands including phospholipids, activated platelets, C-reactive protein and heat shock proteins^60^. Hence, further work is needed to dissect which OLR1 ligands are most pertinent in CLD pathogenesis and how they might be selectively inhibited. A more exhaustive characterisation of the functional impacts of OLR1 inhibition on the macrophage secretome and differentiation will also provide important further insights into the anti-fibrotic effects of targeting SAMac OLR1.

While our focus has been on SAMac heterogeneity and biology, we anticipate that our annotated human liver single-cell atlas will also serve as a valuable resource for researchers to study different aspects of CLD pathogenesis and identify other therapeutic targets. Our analyses have already enabled the distinction of rare hepatic cell types (e.g. neurons, mast cells, basophils, neutrophils and HPCs) and identified several alterations in liver cell abundance in CLD, including expansion of specific subpopulations of endothelial and mesenchymal cells, changes in T cell composition and an accumulation of B and plasma cells. These findings can now be followed up more mechanistically to identify candidate approaches to modulate pathogenic populations in different etiologies of CLD. Furthermore, our fully annotated atlas will support more refined downstream analyses such as reference-based annotation, deconvolution of bulk RNAseq data and integration with spatial transcriptomics. It should be noted that our atlas is based on scRNAseq data, enabling detailed analysis of immune cells. However, other liver cell types such as hepatocytes and HSCs are better captured using single-nucleus RNAseq (snRNAseq)^38,42,61–63^. Moving beyond these studies, future atlases should incorporate both scRNAseq and snRNAseq data from patients at different CLD stages to fully dissect the cellular heterogeneity of immune and non-immune cells.

In conclusion, by generating a fully annotated scRNAseq atlas from healthy and CLD human livers we have resolved previously unknown heterogeneity in SAMacs and identified OLR1 as a new target for liver inflammation and fibrosis. This work forms the basis for a more refined approach to modulating SAMacs and provides a resource for the identification of additional therapeutic strategies in CLD.

## Methods

### Study participants

Local approval for procuring blood samples from CLD patients for *in vitro* analyses was obtained from the Lothian NRS BioResource and Tissue Governance Unit (study number SR574), following review at the East of Scotland Research Ethics Service (reference 15/ES/0094). All subjects provided written informed consent. For histological assessment of MASLD biopsies, anonymized unstained formalin-fixed paraffin-embedded liver biopsy sections were provided by the Lothian NRS Human Annotated Bioresource (study number SR848) under authority from the East of Scotland Research Ethics Service REC 1, reference 15/ES/0094. Unified transparent approval for the pan-Scotland SteatoSITE project2 was provided by the West of Scotland Research Ethics Committee 4 (Reference: 20/WS/0002), Public Benefit and Privacy Panel for Health and Social Care (PBPP; Reference: 1819-0091), Institutional Research & Development departments and Caldicott Guardians.

### Mice

Adult male C57BL/6JCrl mice aged 8–10 weeks were purchased from Charles River. Mice were housed under specific pathogen-free conditions at the University of Edinburgh. All experimental protocols were approved by the University of Edinburgh Animal Welfare and Ethics Board in accordance with UK Home Office Legislation, following ethical guidelines. To induce liver fibrosis, carbon tetrachloride (CCl_4_) (Sigma-Aldrich, 319961) was diluted 1:3 in olive oil (Sigma-Aldrich, 01514) and injected intraperitoneally twice a week for 4 weeks at a dose of 0.4 μl kg^-1^, as previously described^18^. Mice were randomly assigned to receive CCl_4_ or to serve as healthy controls. No sample size calculation or blinding was performed. For scRNAseq, blood and liver tissue were obtained 24 hours, 96 hours and 4 weeks after the final CCl_4_ injection. For scATACseq, blood and liver tissue were obtained 24 hours or 4 weeks following the final CCl_4_ injection. For flow cytometry, liver tissue only was obtained 24 hours, 96 hours, 168 hours and 4 weeks after the final CCL_4_ injection.

For the generation of bone marrow chimera mice, Busilvex (Pierre Fabre, 728981) was diluted to 1 mg/ml in sterile PBS and injected intraperitoneally to a final dose of 10 mg/kg in three-month-old male C57BL/6JCrl mice (CD45.2/CD45.2) over four consecutive days. On day 5, bone marrow cells were isolated from LY5.1 male mice (CD45.1/CD45.2), according to a previously published method^64^. Cells were then enriched for hematopoietic stem cells using the EasySep™ Mouse Hematopoietic Progenitor Cell Isolation Kit (STEMCELL Technologies, 19856), according to manufacturer’s instructions. Next, 8 x 10^5^ enriched cells were injected intravenously into the conditioned C57BL/6JCrl mice. Mice were allowed to recover for 6 weeks prior to CCl_4_ administration

### Preparation of liver and blood single-cell suspensions

For flow cytometry and FACS, mouse liver single-cell suspensions were prepared as previously described, with minor modifications ^65^. In brief, livers were perfused with ice cold PBS via the Inferior Vena Cava (IVC) and then excised and placed in RPMI (Gibco, 21875034). The livers were mechanically chopped into small fragments with a razor blade, followed by digestion in RPMI and an enzyme cocktail containing 0.625 mg ml^−1^ collagenase D (Roche, 11088882001), 0.85 mg ml^−1^ collagenase V (Sigma-Aldrich, C9263-1G), 1 mg ml^−1^ dispase (Gibco, Invitrogen, 17105-041) and 30 U ml^−1^ DNase (Roche, 10104159001) at 37°C for 20 min with agitation. Next, the suspension cells were strained through 100μm filters, centrifuged (300g 5 min) and incubated for 3 minutes in 10% red blood cell (RBC) lysis buffer (BioLegend, 420301). After centrifugation (300g 5 min), cells were resuspended in PEB buffer (PBS containing 0.05% FBS and 0.5 mM EDTA) and passed through a 35 μm filter, prior to blocking and antibody staining. Mouse peripheral blood (100-200μL) was collected from the inferior vena cava into tubes containing a 0.5 mM EDTA solution. Blood was incubated twice in 10% RBC lysis buffer (BioLegend, 420301) for 5 minutes on ice. Cells were then centrifuged (300g 5 min), washed in PEB buffer and filtered through a 35 µm filter, prior to blocking and antibody staining.

### Flow Cytometry and FACS sorting

For liver cell flow cytometry, 2 x 10^6^ liver suspension cells were blocked with anti-mouse CD16/32 antibody (BioLegend, 101302; 1:100) and 10% mouse serum (Sigma-Aldrich, M5905) for 10 minutes at 4°C. Cells were then incubated with primary antibodies for 20 minutes at 4°C. All antibodies, catalogue numbers and dilutions used are listed in supplementary table 16. Cells were then centrifuged (300G 5 min), washed in PEB buffer and incubated with streptavidin BV650 (BioLegend 405232; 1:100) for 20 at 4°C. Flow cytometry compensations were set up using single stained UltraComp eBeads (Thermo Fisher Scientific, 01-2222-42). Controls for gating included ‘fluorescence-minus-one’ (FMO) samples. DAPI (1:1000) and counting beads (Biolegend, 424902; 50ul) were added immediately before sample acquisition.

For intracellular flow cytometry of IL-1β, 5 x 10^6^ isolated liver cells were incubated with 1ul/ml Momensin (BD Bioscience, 554724) in RPMI (Gibco, 21875034) containing 10% FBS (Thermo Fisher Scientific, 10500-064) for 3 hours at 37°C. Control samples were incubated for 3 hours at 4°C. Following incubations, cells were centrifuged (500g, 5 min) and stained with Zombie NIR viability dye (BioLegend, 423105, 1:1000). Thereafter, cells were washed with PEB buffer. Blocking and primary antibody staining were then performed as described above. All antibodies, catalogue numbers and dilutions used are in supplementary table 16. After surface antibody staining, cells were stained with streptavidin-BV421 (BioLegend 405225; 1:100) for 20 minutes at 4°C. Cells were then washed twice in PEB buffer before overnight incubation in fixation buffer (Invitrogen 00-5521-00) at 4°C. After fixation, cells were washed twice with permeabilisation buffer (Invitrogen, 00-8333-56) (1000g, 3 min). Next, cells were incubated with the IL-1β antibody (Supplementary table 16) in permeabilisation buffer for 60 minutes at 4°C. Cells were washed again twice with permeabilisation buffer (Invitrogen, 00-8333-56) (1000g, 3 min) and then resuspended in PEB buffer, before acquisition.

For peripheral blood flow cytometry, blood samples were blocked in anti-mouse CD16/32 antibody (1:100; BioLegend, 101302) and 10% normal mouse serum (Sigma, M5905) for 10 min at 4°C. Cells were then stained using antibodies described in supplementary table 16 for for 20 minutes at 4°C, washed twice in PEB buffer then resuspended in PEB buffer for analysis. DAPI (1:1000) was added immediately before sample acquisition.

All samples were acquired on a six-laser BD LSRFortessa™ Cell Analyzer (BD Biosciences). Data were analysed using FlowJo™ v10.8 (BD Biosciences) Software.

The gating strategies used for identifying liver macrophage subpopulations are shown in Extended Data Fig. 9 and Figure 4G. For flow cytometric analysis of blood from bone marrow chimera mice, viable (DAPI^-^) single CD45.2+ cells were gated. Monocytes were identified as CD11b^+^ CD115^+^ cells, having excluded neutrophils (CD11b^+^ Ly-6G^+^), eosinophils (Siglec F^+^) and lineage+ (NK1.1, Siglec H, CD19, CD3) cells. The degree of blood monocyte donor chimerism for each mouse was calculated as the proportion of monocytes which were CD45.1+, and this value was used to normalise the degree of donor chimerism for each liver macrophage subpopulation from the same mouse.

FACS sorting for mouse scRNAseq and scATACseq was performed as described previously^14^. Following blocking, the isolated liver and blood cells were incubated with primary antibodies for 20 minutes at 4°C. All antibodies, conjugates, dilutions, and their catalogue numbers used for FACS of hepatic NPCs and blood leukocytes are presented in supplementary table 16. After antibody staining, cells were washed with PEB buffer and centrifuged. DAPI viability staining (1:1000) was performed immediately before acquiring the samples on a BD FACSAria^TM^ II (Becton Dickinson). Viable hepatic CD45^+^ cells and peripheral blood CD45^+^ Ly-6G^-^leukocytes were sorted from healthy and CCl_4_-treated mice at stated timepoints following the final CCl_4_ injection. To minimise interindividual variability, cells from 3 mice were pooled for each scRNAseq and scATACseq sample.

### Droplet-based scRNAseq and scATACseq

ScRNAseq was performed using the Chromium^TM^ Single Cell 3′ Library and Gel Bead Kit v2 (10X Genomics, PN-120237) and the Chromium^TM^ Single Cell A Chip Kit (10X Genomics, PN-120236) on the Chromium^TM^ Single Cell Platform (10X Genomics), according to manufacturer’s protocol. After FACS sorting, cells were washed and counted using the Bio-Rad TC20 cell counter (Bio-Rad). Approximately 12,000 cells were loaded on each lane of the chip. For scATACseq, after FACS sorting, cells were washed and nuclei isolated as described in the 10X Genomics demonstrated protocol (document number CG000169), including modifications for low cell input nuclei isolation. Nuclei were counted using a Countess II automated cell counter and processed for scATACseq using the Chromium™ Single Cell ATAC Library & Gel Bead Kit (10X Genomics, PN-1000111) and the Chromium™ Single Cell ATAC Chip E Kit (10X Genomics, PN-1000086), according to the manufacturers protocol. A nuclei recovery of 7000 per lane was targeted. All libraries were sequenced on an Illumina HiSeq 4000.

### Immunofluorescence staining

Immunofluorescence staining of human liver was performed on 5 µm paraffin-embedded tissue sections. After rehydration, heat-mediated antigen retrieval was carried out in separate runs: using pH 9 Tris-EDTA buffer (15 minutes) for the primary antibody for OLR1, and pH 6 sodium citrate buffer (15 minutes) for the primary antibody for GPNMB. Sections were then incubated in 3% hydrogen peroxide (Sigma-Aldrich, H1009) for 10 minutes, washed with PBS containing 0.1% Triton-X (PBS-T), and blocked with protein block (Abcam, ab64226) for 30 minutes. Primary antibodies, diluted in antibody diluent (Abcam, ab64211), were incubated overnight at 4 °C. All antibodies are listed in Supplementary Table 16. The following day, sections were washed with PBS-T and incubated with ImmPRESS HRP Polymer Detection Reagents (Vector Laboratories, MP-7402) for 30 minutes at room temperature (RT). After washing with PBS, initial staining was detected using Cy5 tyramide (Perkin-Elmer, NEL741B001KT) for OLR1 and Cy3 tyramide for GPNMB both at a 1:1,000 dilution. Slides were then washed with PBS-T, followed by an additional round of heat-mediated antigen retrieval using pH 6 sodium citrate buffer (15 minutes) for the second antibody CD68. After PBS washes and protein blocking, sections were incubated with CD68 for 1 hour at RT. Detection was performed using ImmPRESS Polymer and fluorescein-conjugated tyramide, as described above. This staining sequence was repeated for a third round using the antibodies COL1A1 or OLR1. Heat retrieval was performed with pH 6 sodium citrate buffer (15 minutes) for COL1A1 and with pH 9 Tris-EDTA buffer (15 minutes) for OLR1. Both antibodies were incubated overnight at 4 °C, followed by detection with Cy3 tyramide for COL1A1 and Cy5 tyramide for OLR1. After the final staining cycle, all sections were counterstained with DAPI (Sigma-Aldrich, D3571; 1:1,000) and mounted using ProLong Gold antifade reagent (Thermo Fisher Scientific, P36930). Fluorescent images of whole tissue sections were captured using the AxioScan.Z1 slide scanner (Zeiss).

Immunofluorescence staining of stem cell-derived 3D multilineage liver organoids was performed on 4 µm paraffin-embedded tissue sections. After rehydration, heat-mediated antigen retrieval was carried out in Tris-EDTA for 10 minutes. Sections were blocked and stained with primary antibodies as described above. The day after, sections ware stained with fluorescent secondary antibodies for 2 hours. All antibodies, catalogue numbers, lot numbers and dilutions used are in supplementary table 16. Sections were DAPI stained and mounted as described above. Images were taken using an Olympus (Olympus Global) microscope. All images were processed and analysed on Zen Blue (Zeiss) software. Thresholds for all the fluorescent channels were set against the negative controls.

### Cell culture

THP1 human monocytes were cultured in RPMI (Gibco, 31870074) containing 1% Pen-strep (Gibco, 11548876), 1% L-glutamine (Gibco, 25030081) and 10% fetal bovine serum (FBS) (Sigma-Aldrich, F9665). THP1 monocytes were seeded at a density of 1.14 x 10^5^ cells/cm^2^ and differentiated into macrophages after treatment with 100 nM PMA (Sigma-Aldrich, P1585).

For human primary peripheral blood mononuclear cells (PBMC) isolation, SepMate™-50 IVD (STEMCELL Technologies, 85450) kit was used, according to manufacturer’s instructions. In brief, 15 mL of Lymphoprep™ (STEMCELL Technologies, 07801) was added to a SepMate™ tube. An equal volume of patients’ blood was diluted in PBS containing 2% FBS (Gibco, A5256701) and added to the SepMate™ tube. After centrifugation (1200 g, 15 min, RT), the supernatant was collected and washed with PBS with 2% FBS. After centrifugation (300 g, 8 min, 4°C), the PBMC pellet was resuspend in PBS with 2% FBS.

For the enrichment of human primary monocytes, EasySep™ Human Monocyte Enrichment Kit (STEMCELL Technologies, 19059) was used, following manufacturer’s instructions. In brief, tetrameric antibody complexes enrichment cocktail was added to the PBMC suspension and incubated for 5 min at RT. Next, magnetic particles were mixed with the sample and incubated for 5 min at RT. The tube was inserted into EasyEights™ EasySep™ magnet (STEMCELL Technologies, 18103) and incubated for 2.5 min at RT. Primary monocytes were negatively isolated and cryopreserved using CryoStor CS10 (STEMCELL Technologies, 100-1061) in liquid nitrogen. For differentiation of patient-derived monocytes into macrophages, cells were thawed and seeded at a density of 3.13 x 105 cells/cm^2^ in the same media as THP1 cells with the addition of 10 mM HEPES (Gibco, 15630080) and 50 ng/mL human M-CSF (PETROTECH, 300-25).

LX2 human hepatic stellate cells were cultured in DMEM high-glucose with 1% L-glutamine (Sigma-Aldrich, D5796) containing 1% Pen-strep (Gibco, 11548876), and 10% fetal bovine serum (FBS) (Sigma-Aldrich, F9665).

Cell lines and primary cells were cultured in incubators at 37 °C with 5% CO_2_ levels and humidified conditions.

### OLR1 knockdown/inhibition in macrophages

For gene knockdown in THP1 cells, OLR1 Silencer™ Select Validated siRNA (Ambion, 4390824) or Silencer™ Select Negative Control No. 1 siRNA (Ambion, 4390843) was delivered via TransIT-X2^TM^ Dynamic Delivery System (Mirus, MIR 6003) 4 days after PMA-induced differentiation, according to the manufacturer’s manual. Media was changed the day after, and RNA/supernatant collected on day 7. Gene knockdown in patient MDMs was performed 5 days after differentiation of primary monocytes, as described above. RNA/supernatants were collected on day 8. *OLR1* knockdown efficiency was assessed by RT–qPCR. OLR1 inhibition in patient-derived MDMs was performed by adding the human OLR1 Blocking Antibody Golocdacimab/MEDI6570 (MedChemExpress, HY-P99646) or human IgG1 lambda1 Isotype Antibody (MedChemExpress, HY-P99992) for 48 hours before supernatant collection. All supernatants were centrifuged (1000 g, 10 min) to remove debris and used subsequently for ELISA or conditioned media stimulation assays.

### Hepatic stellate cell stimulation assay

For conditioned media experiments, LX2 cells were seeded at a density of 3.13 x 10^4^ cells/cm^2^. The following day, cells were serum-starved for 24 hours. On day 3, the supernatant from primary macrophages was mixed with serum-free media at a ratio of 1/4 and used to treat LX2 cells. On day 4, RNA extraction or proliferation assays were performed.

### Stem cell-derived 3D multilineage liver organoid

Human H9 Embryonic Stem Cells (p46) were used in collaboration with Prof. David Hay’s Lab. To generate 3D multilineage liver organoids, pluripotent stem cells were differentiated into hepatocytes, hepatic stellate cells and endothelial cells and aggregated at a ratio of 10: 3: 1 (cell number 3000:900:300) in 50uL per microwell in Gri3D 96-well plates (Sun Bioscience, Gri3D-96P-S-96-500), according to a previously published protocol^56,66,67^. Human primary monocytes were added at a ratio of 5 (cell number 1500) in 50uL per microwell to spheroids 1 day after cell aggregation. CSF-1 (Peprotech, 300-25-10UG) prepared at a stock concentration of 10ug/mL in PBS and used at 50ng/mL, was added to the spheroid culture media and 150uL media changed was carried out every 48 hours. For optional drug treatment, Pexidartinib hydrochloride (PLX-3397, MedChemExpress) or A83-01 (Tocris Bioscience, 2939/10) were prepared at a stock concentration of 5mM in DMSO and they used at final concentration of 1 uM in spheroid culture media between day 7 and 14 post-monocyte addition. Vehicle (DMSO only) was used as a control. Spheroids were harvested for RNA extraction or immunofluorescence staining.

### RNA extraction and RT-qPCR

Cell lysis and RNA extraction were performed using PureLink^TM^ RNA Mini Kit (Invitrogen, 12183018A) or RNAqueous™-Micro Total RNA Isolation Kit (Thermo Fisher Scientific, AM1931), according to manufacturer’s instructions. cDNA was synthesized using SuperScript™ IV Reverse Transcriptase (Invitrogen, 1809005) following manufacturer’s manual. Quantitative PCR was conducted using LightCycler™ 480 SYBR Green I Master (Roche, 04707516001) either on QuantStudio™ 5 Real-Time PCR System (Applied Biosystems) or LightCycler™ 480 Instrument II (Roche). Primer’s list is in supplementary table 17. The 2^−ΔΔCt^ quantification method was used to analyse gene mRNA expression. Β-ACTIN or GAPDH were used as housekeeping genes for normalisation. Sample gene expression was calculated relative to the average mRNA expression of control groups.

### ELISA

For cytokine expression quantification, LEGENDplex™ assay (BioLegend, 740809) was performed on the supernatant of THP1 cells, according to manufacturer’s instructions. Fluorescence was acquired on a NovoCyte flow cytometer (Agilent). Data were analysed using the BioLegend’s LEGENDplex™ (BioLegend) data analysis software.

### MTT proliferation assay

To assess cell viability and proliferation, the Cell Proliferation Kit I MTT (Roche, 11465007001) was used, according to manufacturer’s instructions. Absorbance was acquired on a BioTek Synergy HTX Multimode Reader (Agilent) at 570 nm. Background signal was subtracted to the absorbance of each sample. Cell proliferation was calculated relative to the average absorbance of control groups.

### Single-cell RNA-seq analysis

#### Human data processing, quality control and integration

Human single-cell RNA-seq data from 8 published studies^14,38–40,42,69–71^ were downloaded from the European Nucleotide Archive (ENA) as paired fastq files (210 samples from 81 patients, supplementary table 1). Sequence alignment and initial sample processing was performed using Cell Ranger (v.7.0.0^72^) and the GRCh38 reference human genome (version GRCh38-2020-A downloaded from 10X Genomics). Subsequent processing and analysis were performed in R (v.4.3.1) using the R package Seurat (v.5.0.0) ^68^, and in python (v.3.10.16). All visualisations were produced using Seurat functions in conjunction with the ggplot2 (v.3.5.1; ^73^) and grid (v.4.3.2; R Core Team [2020]) R packages.

Samples were adjusted for ambient RNA contamination using the R package DecontX (v.1.14.2)^74^, and cells failing quality thresholds (mitochondrial gene percentage > 15% or total unique features < 100) were removed. Patients with fewer than 500 cells passing these thresholds were excluded. As a result, all patients from Tamburini et al. were excluded, leaving 77 patients from 7 studies for downstream analysis.

For patients whose genetic sex was not reported in the original publications, sex was inferred based on the expression of 20 Y chromosome-specific genes: *RPS4Y1, DDX3Y, KDM5D, UTY, USP9Y, ZFY, EIF1AY, TSPY1, PRKY, NLGN4Y, AMELY, PCDH11Y, CDY1, CDY2, VCY, BPY2, DAZ, HIST1H2AY, RBMY,* and *SRY*. 50 cells were randomly selected from each patient, and if any of these genes had non-zero expression in the selected cells, the patient was classified as male; otherwise, as female.

Patient disease status was obtained either from the provided metadata or patient characteristics reported in each study. Liver samples from patients annotated as healthy or those with no fibrosis reported on histology were considered healthy controls, whereas those annotated as diseased or with documented fibrosis on histology were classified as CLD. Where available, the aetiologies of CLD were extracted from the original metadata or were inferred from published histological findings. All other patients with unspecified aetiology were annotated as unknown.

To facilitate efficient integrative analysis of all cells from the 77 remaining patients, 1,000 representative cells were first selected from each sample using the Seurat function ‘*SketchData*’ based on a method described by Hie et al^75^. Normalization, scaling, variable feature identification, and principal component analysis (PCA) were performed on the ’sketched’ data using default Seurat methods, followed by integration using Harmony^76^ and clustering using Seurat. Using the sketched integration as reference, the Seurat function ’*ProjectIntegration’* was then used to integrate the entire dataset. Cluster labels from the sketched cells were projected onto the whole dataset using the function ‘*ProjectData*’ from Seurat.

Wilcoxon Rank Sum tests were performed using the ‘*FindAllMarkers*’ function from Seurat, identifying marker genes from each cluster. Clusters were then manually annotated using previously published literature, identifying 11 cell lineages.

#### Clustering, annotation and atlas construction

Cell lineages were subset individually and the previous pre-processing steps repeated (normalization, scaling, variable feature identification, and PCA). As before, cells were clustered and then annotated based on expression of known marker genes. Doublet clusters were identified manually based on their expression of multiple lineage markers and removed. Low-quality clusters (median nFeature < 800) were also removed. After quality control for each lineage, the final atlas included 649,295 cells.

Annotations from each cell lineage (subsequently referred to as ‘level 3’ annotation) were mapped back to the complete atlas. Cluster similarities were assessed using the function ‘*buildSNNGraph*’ from R package scran (v.1.30.2^77^) and the function ‘*pariwiseModularity*’ from R package bluster (v.1.12.0;^78^). Based on the results, clusters from level 3 Annotation were manually merged into 34 grouped clusters, defined as the ‘level 2’ annotation. The same similarity testing and merging procedures were then applied to the level 2 clusters, resulting in the manual identification of 16 ‘level 1’ clusters.

For atlas visualization, a ShinyApp was built using R package ShinyCell (v.2.1.0^79^). In addition, a Loupe file was generated using loupeR (v.1.1.4 https://github.com/10XGenomics/loupeR).

#### Differential abundance and expression testing

Differential composition analysis (DCA) was performed on level 3 annotations of selected lineages using MiloR (v1.8.1)^44^. First, cellular states were modelled as neighbourhoods based on the k-nearest neighbour graph (KNN). Differential neighbourhood abundance testing was then performed comparing disease status (healthy vs CLD).

For the MP lineage, additional differential abundance tests were also performed for each level 3 MP cell type, comparing healthy vs. CLD, male vs. female, and healthy vs. each annotated disease aetiology. Analysis was restricted to patients with more than 50 MPs. The relative abundance of each cell type was calculated for each patient, and relative abundances between groups tested using unpaired two-sample t-tests, implemented via the ‘*stat_compare_means* ‘function from the ggpubr R package.

Differential gene expression analysis (DGEA) was performed comparing the MP clusters, SAMac1 and SAMac2 using the ‘Model-based Analysis of Single-cell Transcriptomics’ (MAST) framework^80^, deployed using the Seurat function ‘*FindMarkers*‘.

#### Cell-cell interaction analysis

Ligand-receptor analysis was performed on the Level 3 atlas using python packages liana-py (v.1.4.0^50^) and cell2cell (v.0.7.4^50^). To accommodate python analysis, the complete integrated dataset was transformed from a seurat object to h5ad format using R packages sceasy (v.0.0.7) and reticulate (v.1.35.0). Analysis was limited to the MP clusters SAMac1 and SAMac2 as senders and level 3 mesenchymal cell clusters as receivers. Ligand-receptor score calculations were iteratively run across individual patient samples using the liana-py function: ‘*rank_aggregate.by_sample’*. Ligand-receptor scores were input to cell2cell as a four-dimensional tensor (sample × sender cell × receiver cell × ligand-receptor pair). Using the Tensor-cell2cell pipeline, tensor decomposition was performed extracting unique context dependent patterns of intracellular communication in the form of 10 factors. Prior to decomposition, elbow analysis was performed to select the optimal number of factors. Ligand-receptor pairs associated with each factor were retrieved using the cell2cell function, ‘*tensor_downstream.get_lr_by_cell_pairs*’.

#### Mouse scRNAseq analysis

Analysis was performed on 12 mouse scRNAseq datasets (n=6 viable CD45+ cells from liver and n=6 viable CD45+ Ly-6G-cells from peripheral blood) in total. Blood and liver cells were obtained from chronic CCl_4_ treated mice at three timepoints following the final CCl_4_ injection (24 hours, 96 hours and 4 weeks) and age-matched uninjured controls.

Sequence alignment and initial sample processing was performing using Cell Ranger (v.3.0.2^72^) and the mm10 reference human genome (version mm10-3.0.0 downloaded from 10X Genomics). Subsequent processing was performed in R (v.4.2.2) using the R package Seurat (v.4.1.3). All visualisations were produced using Seurat functions in conjunction with the ggplot2 (v.3.2.1^73^) and grid (v.3.6.3; R Core Team [2020]) R packages.

Cells failing quality thresholds (mitochondrial gene percentage > 30% or total unique features < 300) were removed. Normalization, scaling, variable feature identification, and PCA were performed using default Seurat methods.

After merging the 12 datasets, dimensional reduction was performed (uniform manifold approximation and projection; UMAP) showing complete overlap of the different samples. Clustering was then performed iteratively, identifying and removing doublet clusters based on their expression of multiple lineage markers. After quality control, the final dataset included 27,070 cells.

Clustering of the final dataset was then performed using default methods, and marker genes for each cluster were identified by Wilcoxon rank sum test, facilitating manual annotation of cell types.

The mononuclear phagocyte lineage (MP) was isolated and re-analysed as described previously (normalization, scaling, variable feature identification, PCA). Clustering was then performed using the first 17 PCs and a resolution of 0.7. As before, annotation was performed manually based on the expression of known marker genes.

Differentially expressed genes between the SAMac1 and SAMac2 clusters were identified using the ‘Model-based Analysis of Single-cell Transcriptomics’ MAST approach^80^. Genes lists were filtered to include only genes that are expressed in >10% of cells with an absolute log-transformed fold change greater than 0.5 and Bonferroni adjusted p-value < 0.01.

#### Mouse scATACseq analysis

Analysis was performed on 8 mouse scRNAseq datasets (n=4 viable CD45+ cells from liver and n=4 viable CD45+ Ly-6G-cells from peripheral blood) in total. Blood and liver cells were obtained from chronic CCl_4_ treated mice at two timepoints following the final CCl_4_ injection (24 hours and 4 weeks) and age-matched uninjured controls.

Sequence alignment and initial sample processing was performed using Cell Ranger ATAC (v.1.1.0^81^) and the mm10 reference mouse genome (version cellranger-atac-mm10-1.1.0 downloaded from 10X Genomics). Gene annotations for the mouse genome (mm10) were retrieved from the EnsDb.Mmusculus.v79 (v.2.99.0) database. Subsequent preprocessing, clustering, annotation, motif discovery, and downstream analysis was performed in R (v.4.2.2) using the R packages Seurat (v.4.1.3) and Signac (v.1.14.0^82^)

Cells failing quality control thresholds were removed (reads in peaks <300 or >50,000; fraction of reads in called peak regions (FRiP) <0.05; blacklist ratio >0.125; transcription start site (TSS) enrichment <2; number of detected features and counts <3000). After quality control, samples were integrated using Harmony (v.1.0.0^71^), creating a single dataset with 29,272 cells.

Normalization and dimension reduction was performed using Latent Semantic Indexing (LSI), as described by Cusanovish et al^83^. Clustering was performed using LSI components 2 to 30 to construct the KNN graph, and a resolution of 3.5, to partition the dataset into 16 final clusters. LSI component 1 was omitted from the analysis due to its correlation with sequencing depth.

#### Cluster Identity with scRNAseq and Peak Calling

Using cross-modality integration and label transfer^84^, scATACseq cells were annotated using the mouse scRNAseq dataset. First, anchors were identified between the query (scATACseq) and reference (scRNAseq) datasets using canonical correlation analysis (CCA), followed by the transfer of annotations from the reference to query dataset.

MPs were subset (14,613 cells) and subsequent processing was performed using R package, ArchR (v.1.0.2^85^) and the precompiled mm10 genome (BSgenome.Mmusculus.UCSC.mm10). Peak calling for MPs was performed using python package MACS2 (v.3.0.3^86^) installed in R via the Anaconda package repository using R package Herper (v.1.18.0^87^)

After subsetting, the top abundant peak features were recalculated and used as input for further normalization and clustering. As before, cells failing quality control thresholds were removed (reads in peaks <300 or >50,000; fraction of reads in called peak regions (FRiP) <0.05; blacklist ratio >0.125; transcription start site (TSS) enrichment <2; number of detected features and counts <3000). TF-IDF normalization was performed, followed by LSI dimension reduction. Samples were then reintegrated using Harmony implemented within ArchR. Gene scores were calculated, providing an estimation of how highly expressed a gene will be based on the accessibility of local regulatory elements. Label transfer was then performed using the MP scRNA-seq dataset as reference.

#### Differential Peaks and TF Motif Enrichment

Cluster specific peaks were identified using the ArchR function ‘*addMarkerFeatures*’ to perform pairwise Wilcoxon Rank Sum tests. By setting the ‘bias’ parameter to adjust for TSS enrichment and total unique fragments per cell, differences in quality between clusters are adjusted for during testing. Differential peaks were also identified between the MP clusters SAMac1 and SAMac2.

Using the chromVARmotifs R package (v.1.30.0^81^), motif sets from the CIS-BP repository^88^ were retrieved. To identify motifs that are enriched in SAMac1 versus SAMac2, differential peaks with false discovery rate (FDR) <= 0.1 and log2 adjusted fold change (log2FC) >= 0.5 were subset and tested for motif enrichment.

### Human Liver bulk RNA-seq analysis

RNA-seq and histology data was generated as described in detail in the original manuscript^2^. For correlation between OLR1 gene expression and time-to-event analysis, biopsy cases with available RNAseq from the SteatoSITE multimodal database^2^ were used (R 4.4.3). Normalised counts for OLR1 were available as part of the dataset^2^. Decompensation events in the clinical data extract were defined by a combination of ICD codes and UK OPCS-4 codes identifying procedures relating to cirrhosis-related hospital admissions activity as previously described^89^. Decompensation event analysis was only undertaken on biopsy cases for which the first decompensation-related coding was present in the clinical data extract after the biopsy date and analysis was undertaken using death as a competing risk. The cutpoint for OLR1 gene activity was calculated using surv_cutpoint() from the ‘survminer’ (0.4.9) package, applying maximally selected rank statistics of the ‘maxstat’ package (0.7.25) with a minimum proportion of 0.25. Kaplan–Meier estimator curves of all-cause mortality or decompensation events for assigned high/low risk groups were compared by log-rank testing in ‘survminer’.

### Statistical Analysis

All non-bioinformatics data were analysed using GraphPad Prism 9 software. All graphs are shown as mean±standard error of mean (SEM). Normality was tested using the Shapiro-Wilk test and subsequent parametric and non-parametric statistical analysis were chosen accordingly. Statistically significant comparisons are highlighted as *=P≤0.05, **=P≤0.01, ***=P≤0.001, ****=P≤0.0001.

## Data Availability

Human scRNAseq data was analysed from the following publicly available datasets: GSE243981^38^, GSE168933^71^, GSE192742^42^, GSE115469^39^, E-MTAB-10143^40^, GSE136103^14^, GSE181483^70^ and GSE129933^69^. An online data browser for annotated human scRNAseq data is avallabile at https://shiny.igc.ed.ac.uk/Human_Liver_scRNAseq_Atlas/. A publicly accessible browser for mouse MP data is available at: https://shiny.igc.ed.ac.uk/mouseMPshinyApp/. Hepatic bulk RNA-seq data from SteatoSITE cohort are deposited in the European Nucleotide Archive (https://www.ebi.ac.uk/ena; study accession number: <u>PRJEB58625</u>). SteatoSITE gene expression data are also freely available for user-friendly interactive browsing online at https://shiny.igc.ed.ac.uk/SteatoSITE_gene_explorer/.

## Code Availability

All code for the scRNAseq and scATACseq analyses are available at https://git.ecdf.ed.ac.uk/ramachandran-lab/olr1-paper. The code used in this study for the bulk RNA-seq data analyses of samples from the SteatoSITE cohort can be made available by contacting the corresponding author (J.A.F.). Access to code will be granted for requests for academic use within four weeks of application. R is required software to use the code.

## Acknowledgments

We thank Professor Stuart J. Forbes for appraising and providing feedback on the manuscript. We thank the patients who donated samples for this study; J. Davidson and J. Black, of the Scottish Liver Transplant Unit for assistance with consenting patients for this study; the staff of the Scottish Liver Transplant Unit, Royal Infirmary of Edinburgh for assistance in procuring human blood samples; W. Mungall and BVS staff for technical support with *in vivo* work. Flow cytometry data was generated with support from the IRR flow cytometry and cell sorting facility, University of Edinburgh. Microscopy and image analysis were carried out at and with the help of the IRR Imaging facility at the University of Edinburgh. P.R. is funded by an MRC Senior Clinical Fellowship (MR/W015919/1). N.C.H. is supported by a Wellcome Trust Senior Research Fellowship in Clinical Science (ref. 219542/Z/19/Z). Construction of the SteatoSITE data commons was funded by Innovate UK (Precision medicine: impacting through innovative technology - Ref: TS/R017581/1). This publication is part of the Human Cell Atlas (www.humancellatlas.org/publications).

## Author Contributions

E.P, K.K. and F.C. contributed equally. E.P., K.K., J.T., J.L., S.V., M.H., J.R.W.-K., G.F., and G.P. performed computational analysis and interpretation of scRNAseq data. T.J.K. and J.A.F. performed computational analysis and interpretation of SteatoSITE data. E.P., F.C., J.T., M.G., P.K.Q., A.A., I.B., E.F.S., performed experimental design, data generation, data analysis and interpretation. A.K., G.R-J., C.C.B. and D.C.H. provided advice and expertise on experimental design and data interpretation. L.J.P, C.A.V., N.C.H., T.J.K. and J.A.F. provided expertise and advice on data analysis and interpretation. E.P, K.K., F.C., J.L, J.T., T.J.K, J.A.F. and P.R. wrote the manuscript. P.R. conceived the study, designed the experiments, interpreted data and supervised the study. Address correspondence to P.R.

## Extended Data Figures

**Extended Data Fig. 1:**
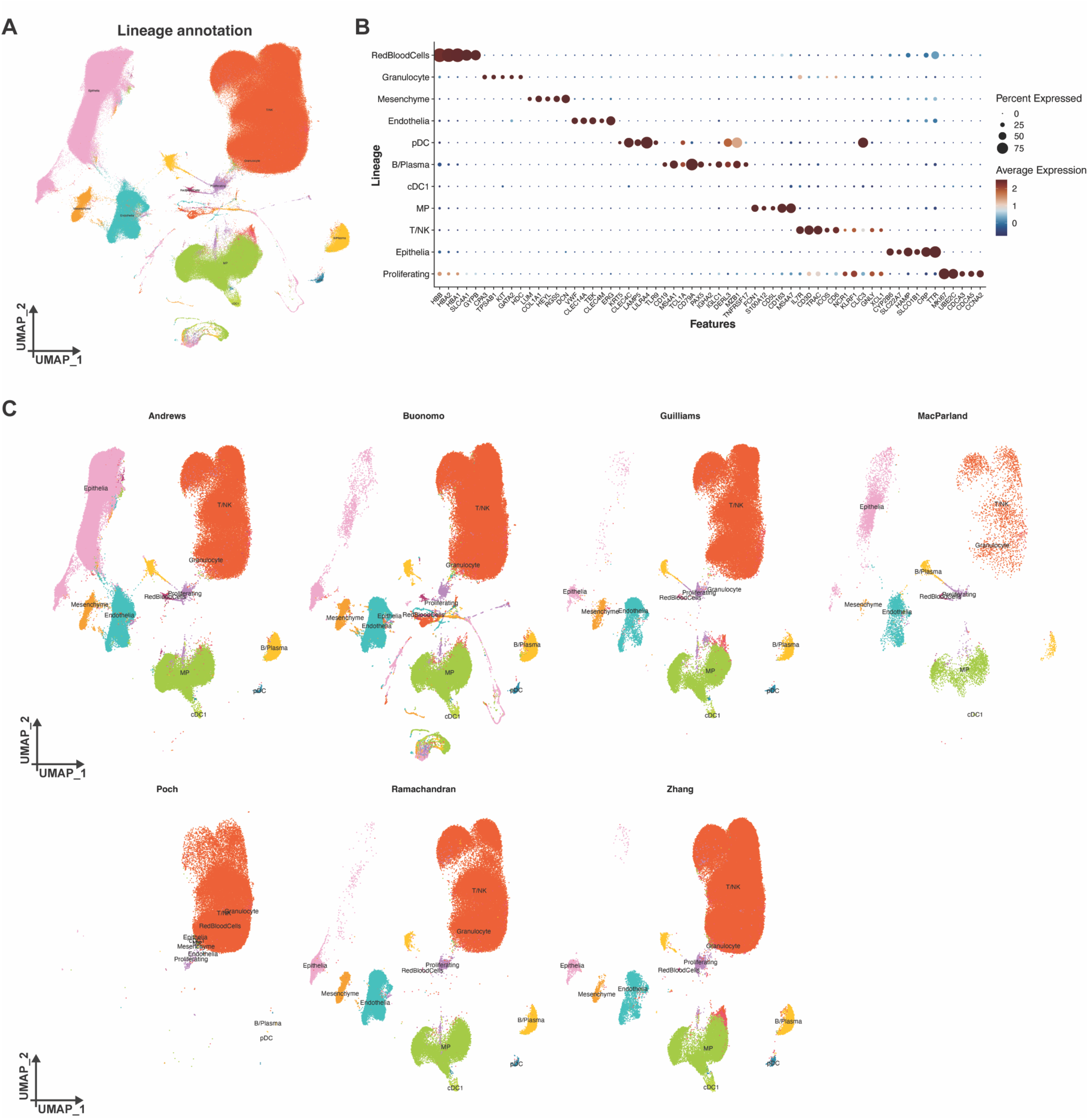
Integration and lineage annotation of human liver scRNAseq data. (**A**) UMAP of integrated scRNAseq data (843,457 cells; 77 patients; 7 studies) coloured and labelled by cell lineage annotation. (**B**) Dotplot showing mean expression of marker genes and percentage of cells expressing them for each lineage cell type. (**C**) UMAP of integrated scRNAseq data split by original study. Coloured and labelled by cell lineage annotation.

**Extended Data Fig. 2:**
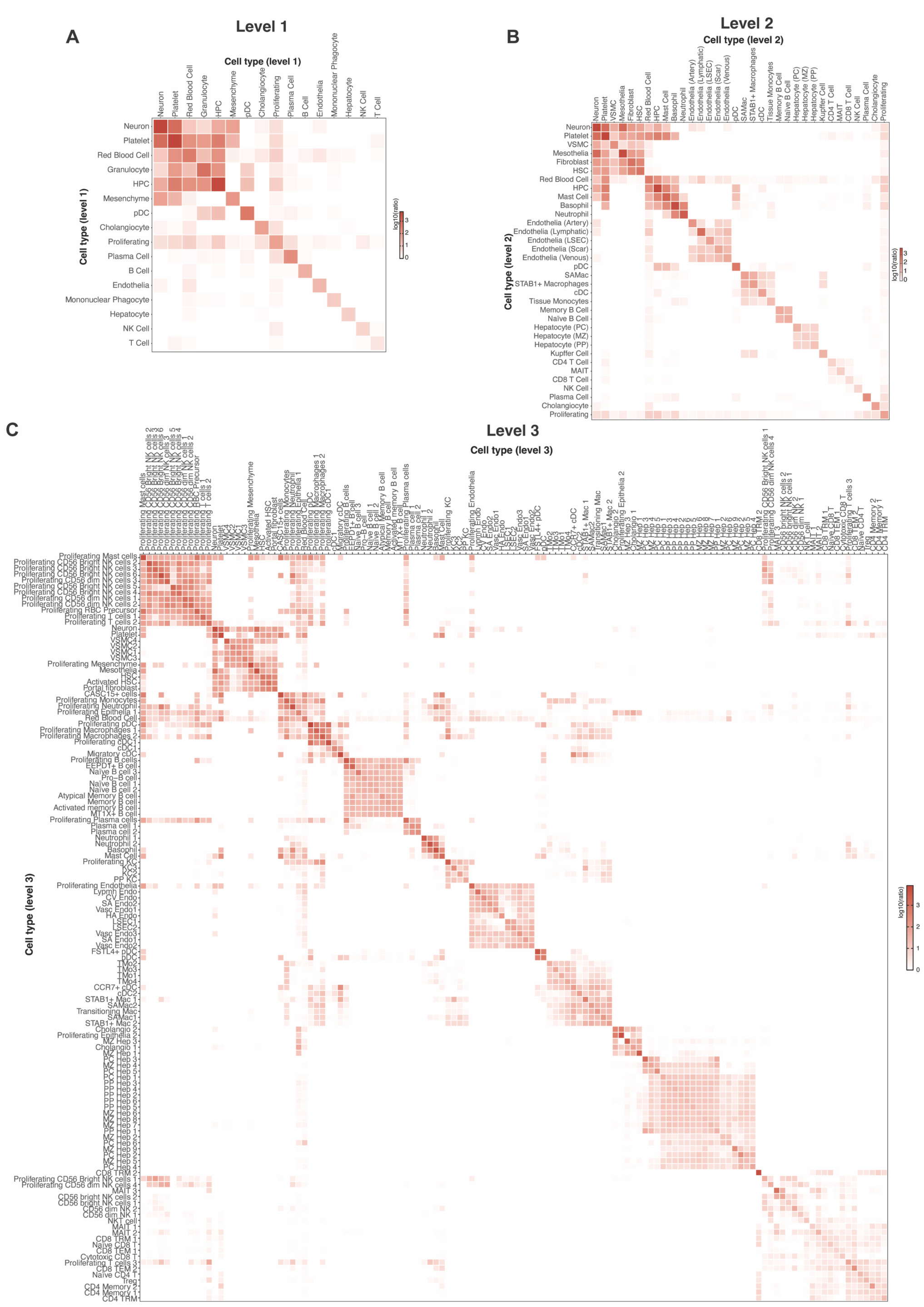
Hierarchical clustering of human liver cell states. (**A**-**C**) Plots of transcriptional similarity between stated cell types. Colour scale indicates log_10_-transformed pairwise modularity ratios between clusters. (**A**) Similarity plot for annotated level 1 cell types in scRNAseq atlas. (**B**) Similarity plot for annotated level 2 cell types in scRNAseq atlas. (**C**) Similarity plot for annotated level 3 cell types in scRNAseq atlas.

**Extended Data Fig. 3:**
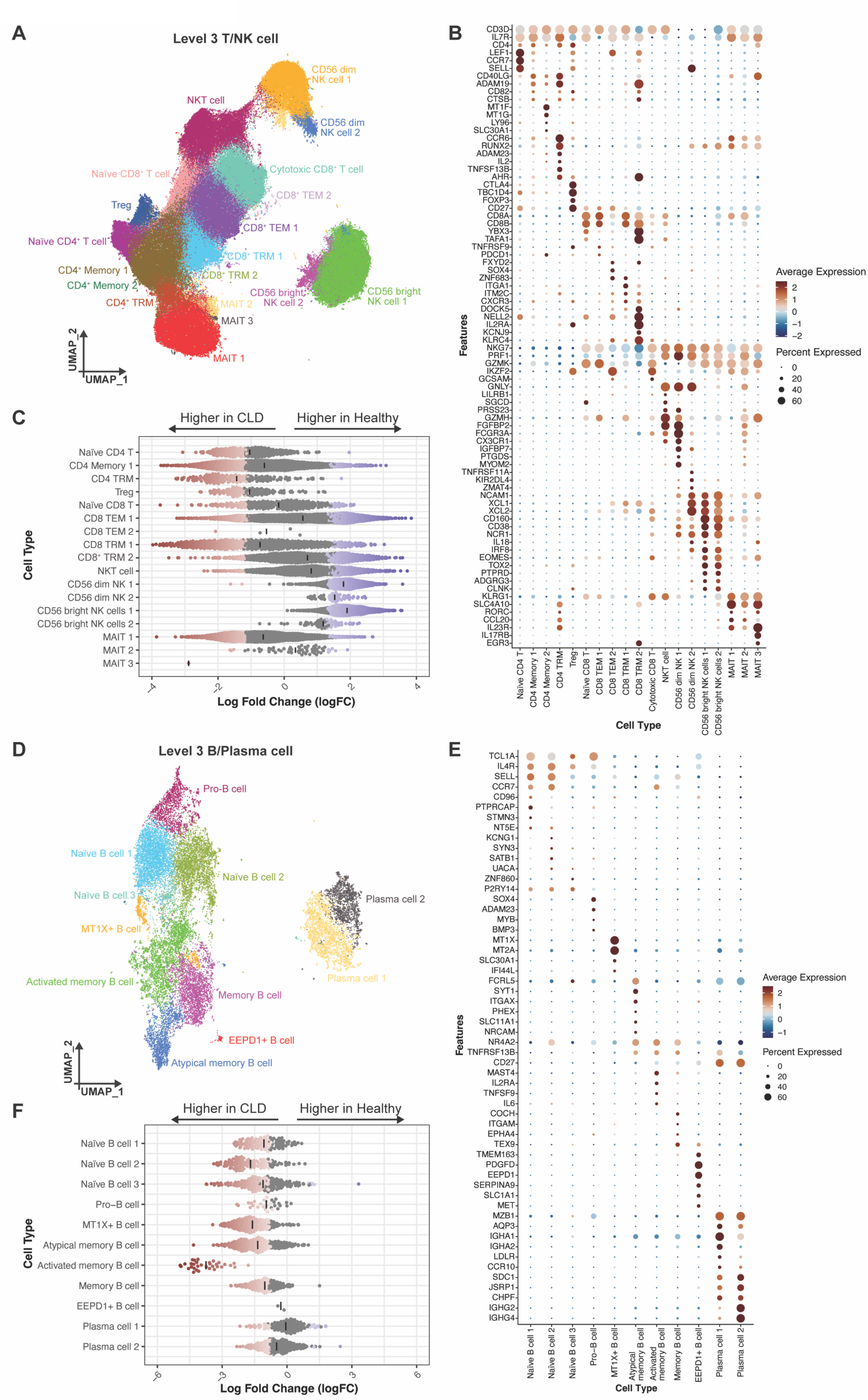
Structure of human liver single cell atlas. Chord diagrams showing hierarchical structure of human liver single-cell atlas. Level 1 annotation represents broad cell lineages which are subdivided into a moderate degree of cell type specificity (level 2 annotation), which are further subdivided into a fine-grained annotation of different cell states (level 3 annotation).

**Extended Data Fig. 4:**
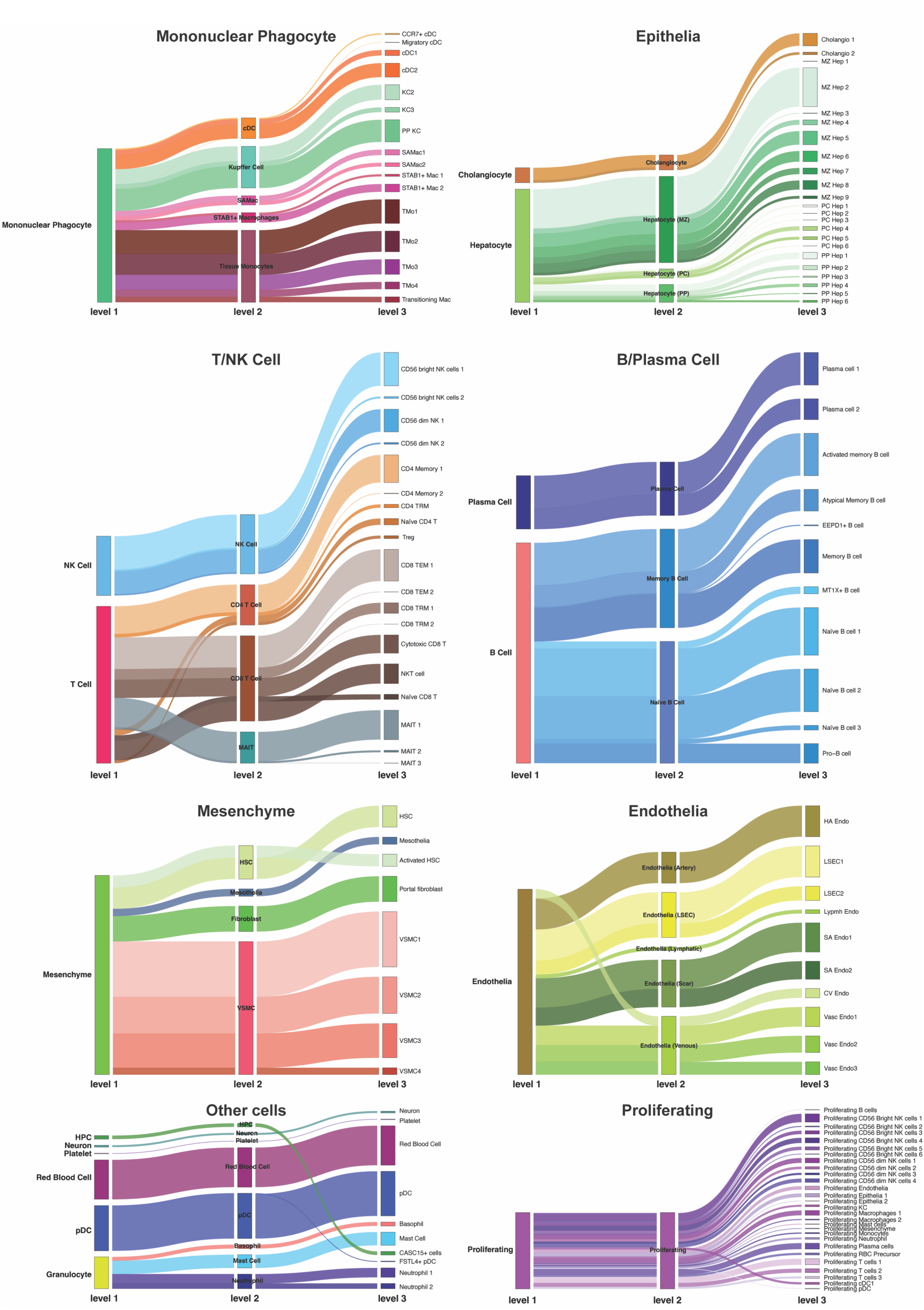
SAMac subpopulations expand in CLD livers. (**A**) UMAPs of mononuclear phagocytes (81,577 cells) coloured and labelled by level 3 annotation. UMAPs split by original study. (**B**) Stacked barplots of proportions of cells of each level 3 MP cell type across the scRNAseq studies. Bars coloured by level 3 MP cell type. Total number of MPs from each study is stated at the top of each bar. (**C**) Differential abundance analysis of each level 3 MP subtype between healthy (blue bars; n=38) and CLD (red bars; n=25 patients across all of the samples using Welsh’s t-tests. \**p*<0.05, \*\**p*<0.01, \*\*\**p*<0.001, \*\*\*\**p*<0.0001, ^ns^*p*≥0.05. (**D**) Differential abundance analysis of each level 3 MP subtype between male (green bars, n=29) and female (yellow bars, n=34) patients across all of the samples using Welsh’s t-tests. \**p*<0.05, ^ns^*p*≥0.05. (**E**) Differential abundance of SAMac1 and SAMac2 between healthy (blue) patients and patients with different aetiologies of CLD (red), using Welsh’s t-tests. Healthy n=38; ALD (alcohol-related liver disease) n=2; MASLD n=4; PBC (primary biliary cholangitis) n=3, PSC (primary sclerosing cholangitis) n=8

**Extended Data Fig. 5:**
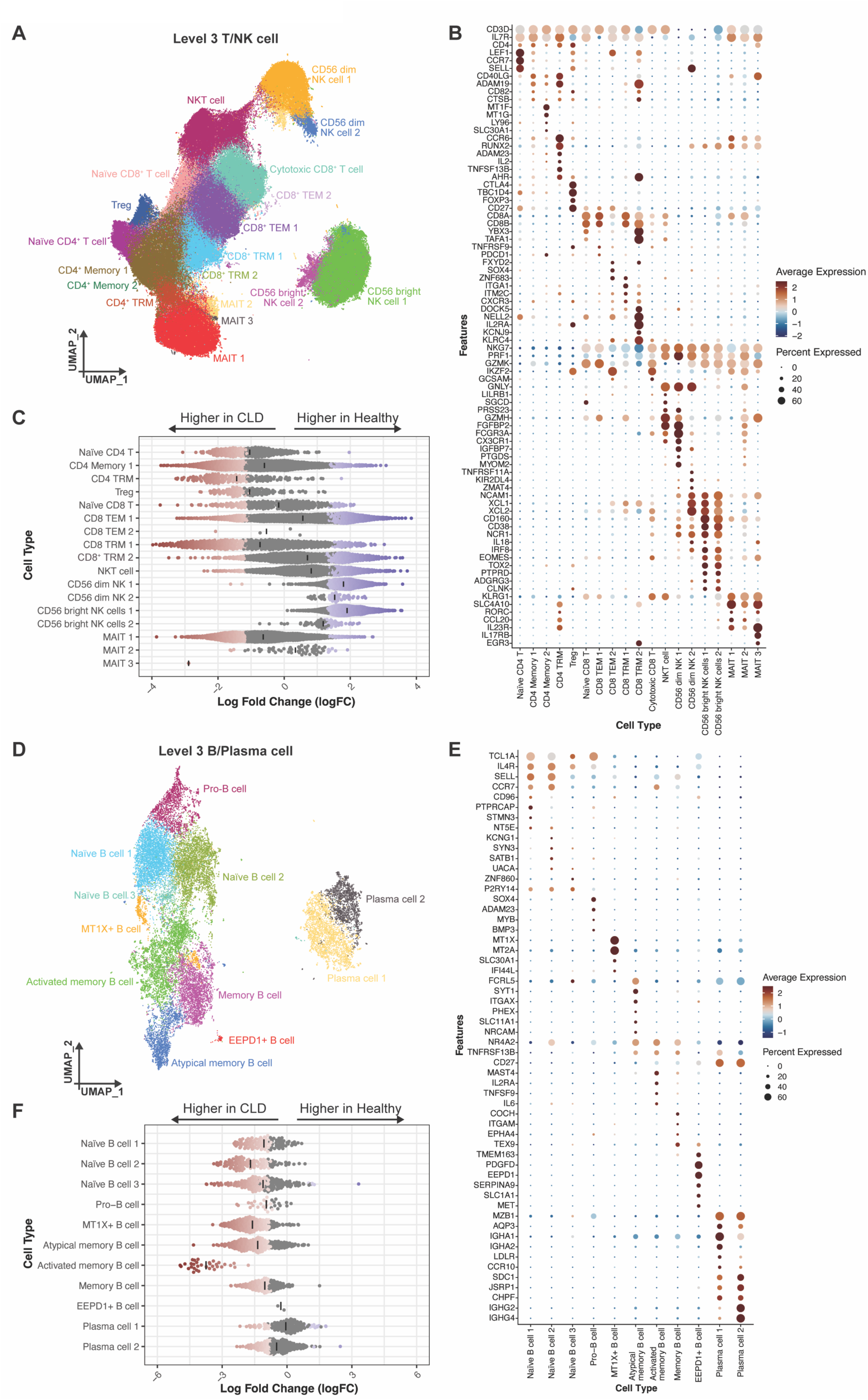
Alterations in hepatic lymphoid cells in CLD patients. (**A**) UMAP of T/NK cells (393,035 cells) coloured and labelled by level 3 annotation. (**B**) Dotplot showing mean expression of marker genes and percentage of cells expressing them for each level 3 T/NK cell type. (**C**) Beeswarm plot of the log fold changes in Milo cell neighbourhoods in T/NK cells by CLD status. Milo neighbourhoods grouped into each level 3 T/NK cell type subcluster. Neighbourhoods with a significant (FDR <0.05) change in cellular abundance are coloured, with blue representing enrichment in healthy and red denoting expansion in CLD. Black lines denote mean logFC for each cell type. (**D**) UMAP of B/Plasma cells (22,854 cells) coloured and labelled by level 3 annotation. (**E**) Dotplot showing mean expression of marker genes and percentage of cells expressing them for each level 3 B/Plasma cell type. (**C**) Beeswarm plot of the log fold changes in Milo cell neighbourhoods in B/Plasma cells by CLD status. Milo neighbourhoods grouped into each level 3 B/Plasma cell type subcluster. Neighbourhoods with a significant (FDR <0.05) change in cellular abundance are coloured, with blue representing enrichment in healthy and red denoting expansion in CLD. Black lines denote mean logFC for each cell type.

**Extended Data Fig. 6:**
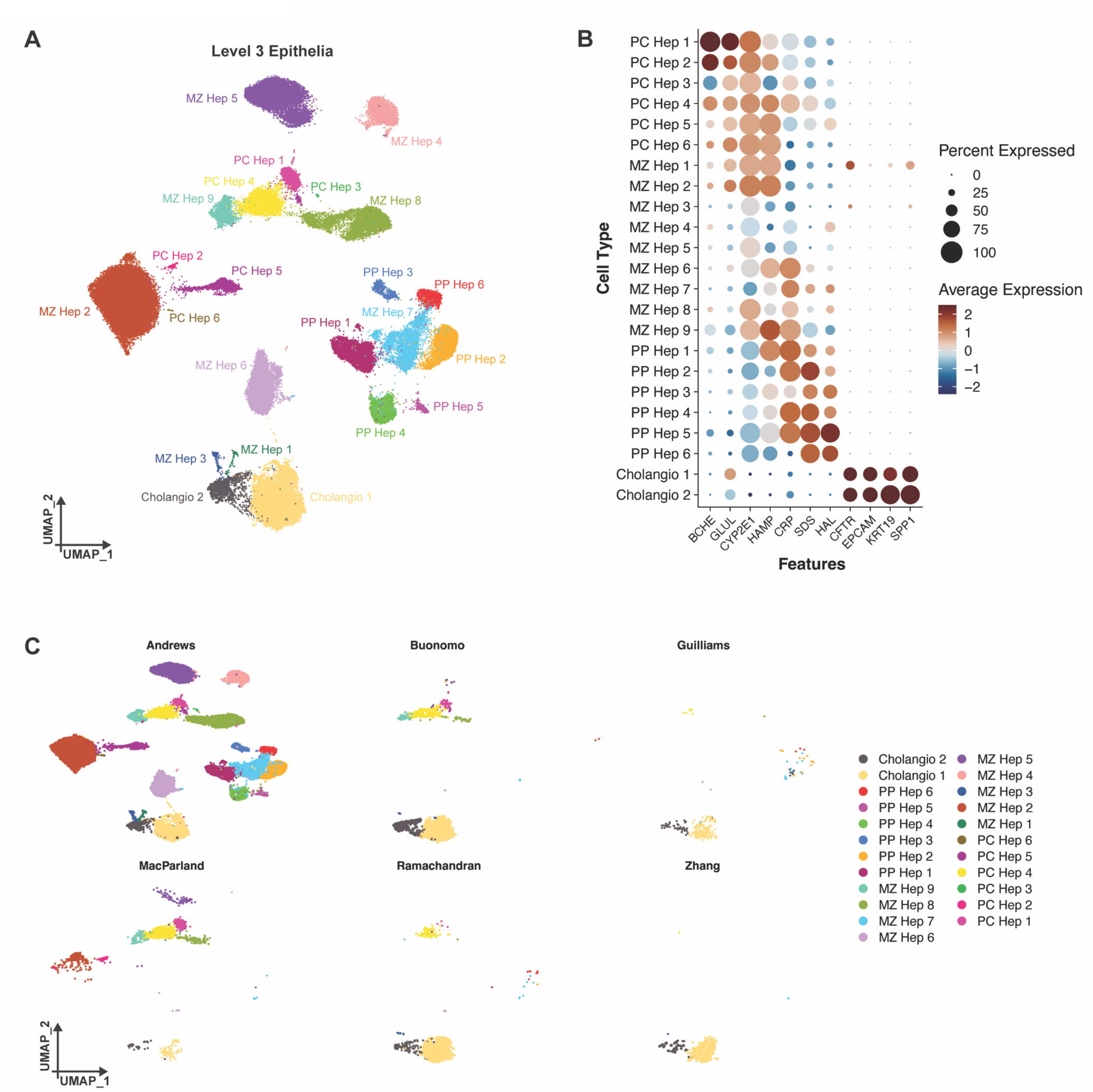
Defining liver epithelial cell heterogeneity. (**A**) UMAP of liver epithelial cells (91,716 cells) coloured and labelled by level 3 annotation. (**B**) Dotplot showing mean expression of marker genes and percentage of cells expressing them for each level 3 epithelial cell type. (**C**) UMAPs of epithelial cells coloured and labelled by level 3 annotation. UMAPs split by original study.

**Extended Data Fig. 7:**
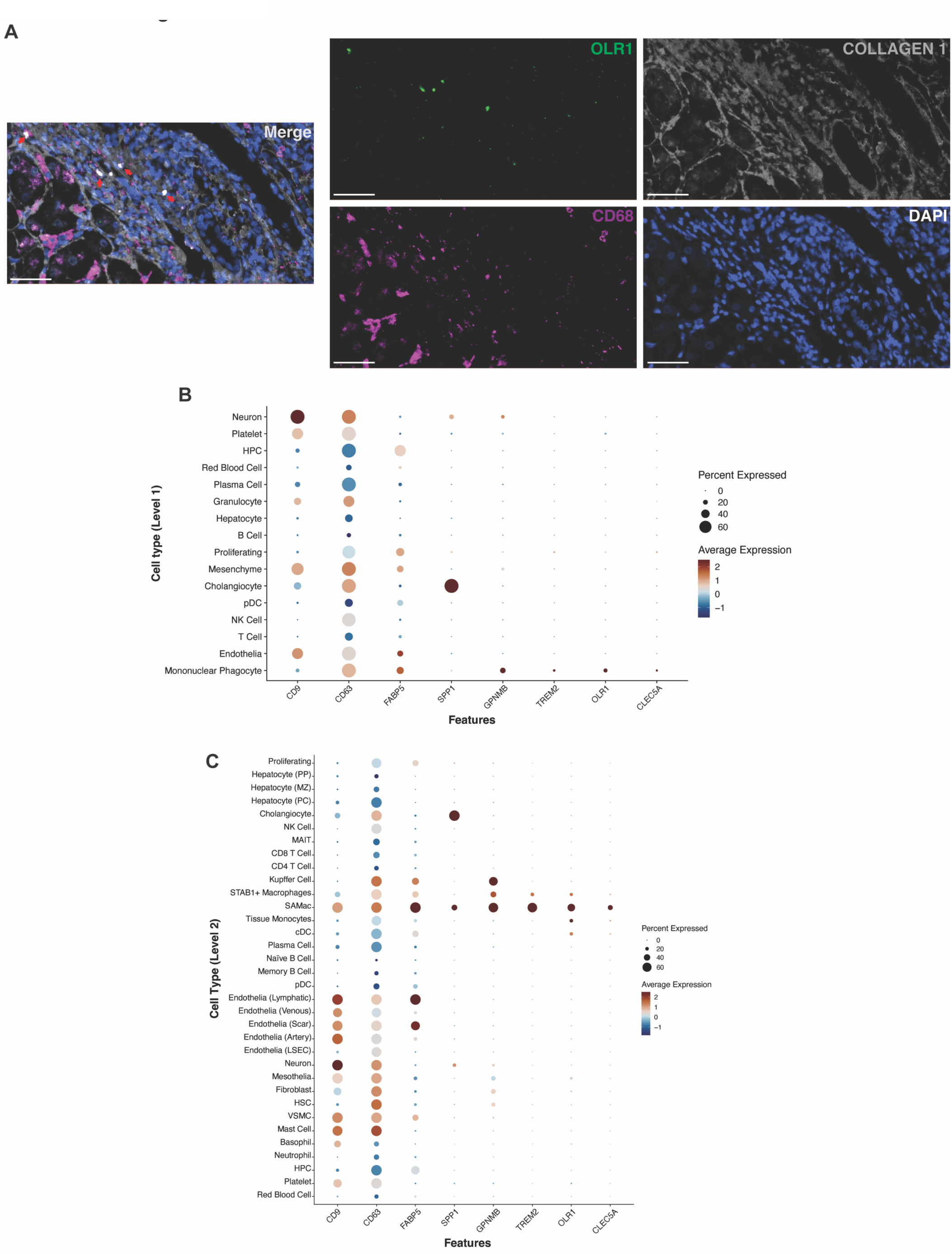
OLR1 expression distinguishes a subpopulation of SAMacs. (**A**) Representative multiplex immunofluorescence image of the fibrotic niche of human CLD liver. Markers are Collagen 1 (grey), CD68 (magenta), OLR1 (green) and DAPI (blue). Scale bar=50μm. Merged and individual fluorescent channels shown. Red arrows indicate OLR1+ CD68+ SAMacs. (**B**) Dotplot showing mean expression of stated genes and percentage of cells expressing them for each level 1 cell type. (**C**) Dotplot showing mean expression of stated genes and percentage of cells expressing them for each level 2 cell type.

**Extended Data Fig. 8:**
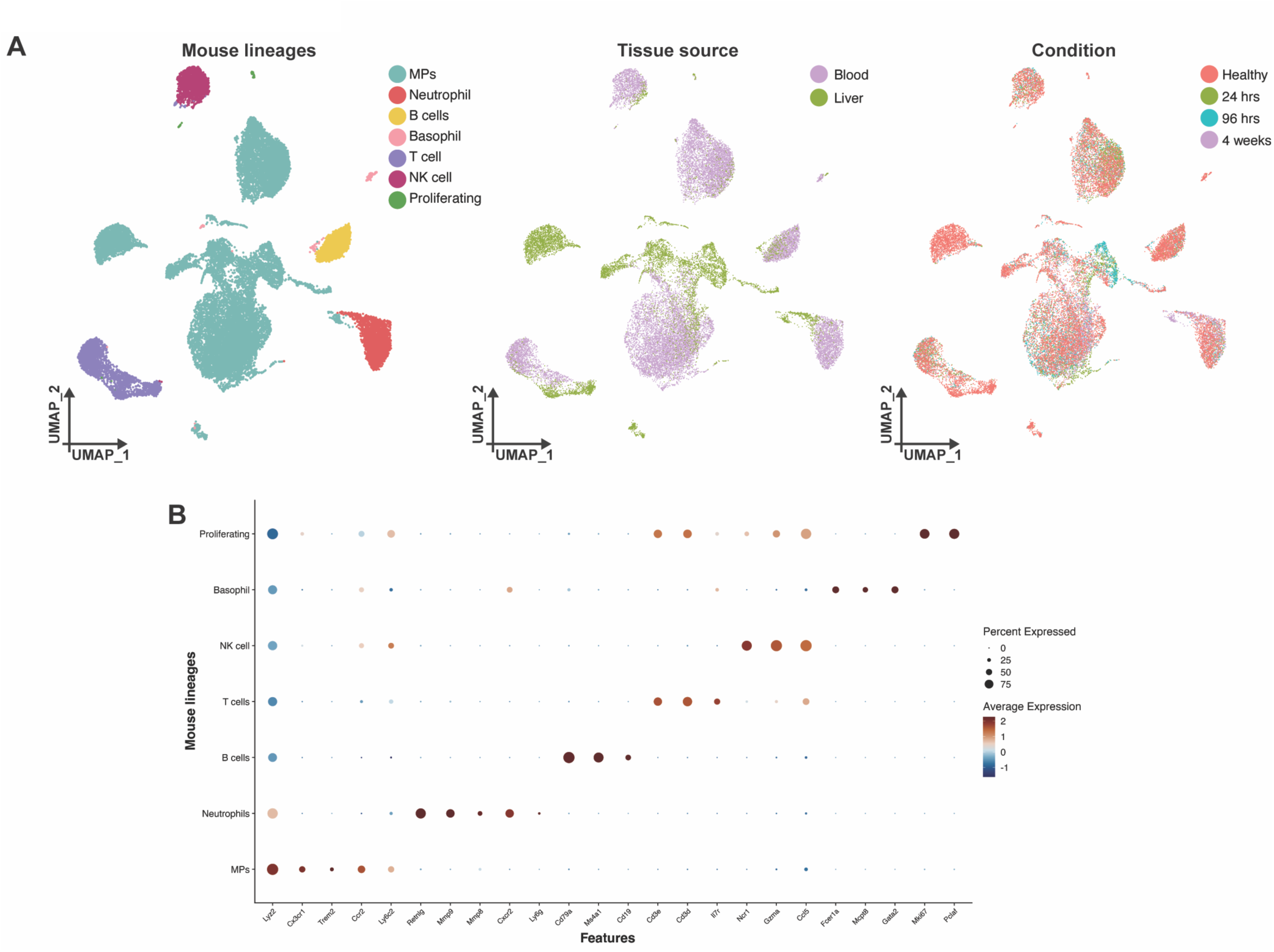
Identifying mouse MPs for subsequent scRNAseq analysis. (**A**) UMAPs of mouse immune cell scRNAseq data (27,070 cells) from blood and liver of chronic CCl_4_ model (see Figure 4A). UMAPs coloured and labelled by cell lineage annotation (left), by tissue source blood vs liver (middle) and by condition/CCl_4_ model timepoint (right). (**B**) Dotplot showing mean expression of marker genes and percentage of cells expressing them for each mouse cell lineage.

**Extended Data Fig. 9:**
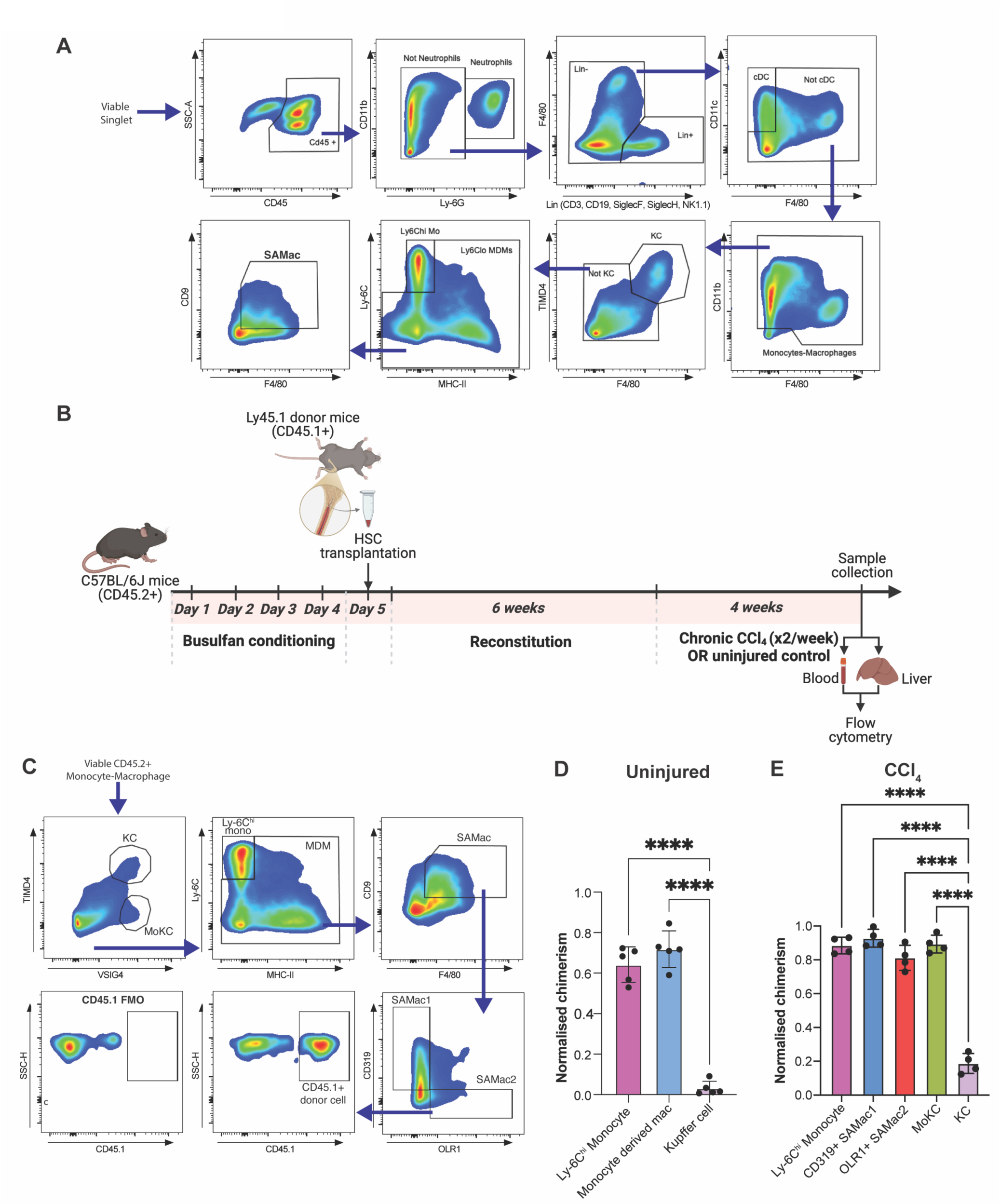
Dissecting SAMac origin and kinetics in mouse CLD. (**A**) Flow cytometry gating strategy for study of SAMac kinetics in mouse liver. Cells were pre-gated as viable (DAPI negative) singlets (based on FSC-A vs FSC-H and SSC-A vs SSC-H). SAMac population is then gated for OLR1 and CD319 expression as shown Fig 4G. (**B**) Overview of experimental workflow for mouse bone marrow chimera CLD model. Wild-type mice (CD45.2+) were used as recipients and CD45.1+ mice as bone marrow donors. Created in BioRender. Colella, F. (2025) https://BioRender.com/fnfo96e. (**C**) Flow cytometry gating strategy for bone marrow chimera studies to quantitate macrophage subpopulation chimerism in mouse liver. Cells were pre-gated as viable (DAPI negative) singlets (based on FSC-A vs FSC-H and SSC-A vs SSC-H) and monocyte-macrophages defined as shown in A. The gated macrophage subpopulations shown were then all assessed for the degree of chimerism (% CD45.1+). MDM=monocyte-derived macrophage, KC=Kupffer cell, moKC=monocyte-derived Kupffer cell (**D**) Bar plot of degree of chimerism of liver macrophage subpopulations, quantitated by flow cytometry of uninjured mice following the busulfan chimera protocol. Bars indicate mean ± S.D. with individual dots representing independent mice. Normalised chimerism calculated for each mouse by normalising to the % of peripheral blood monocytes which are donor-derived (CD45.1+). Ordinary one-way ANOVA with Tukey’s multiple comparisons test. \*\*\*\**p*<0.0001. (**E**) Bar plot of degree of chimerism of liver macrophage subpopulations, quantitated by flow cytometry of chronic CCl_4_ treated mice following the busulfan chimera protocol. Bars indicate mean ± S.D. with individual dots representing independent mice. Normalised chimerism calculated for each mouse by normalising to the % of peripheral blood monocytes which are donor-derived (CD45.1+). Ordinary one-way ANOVA with Tukey’s multiple comparisons test. \*\*\*\**p*<0.0001.

**Extended Data Fig. 10:**
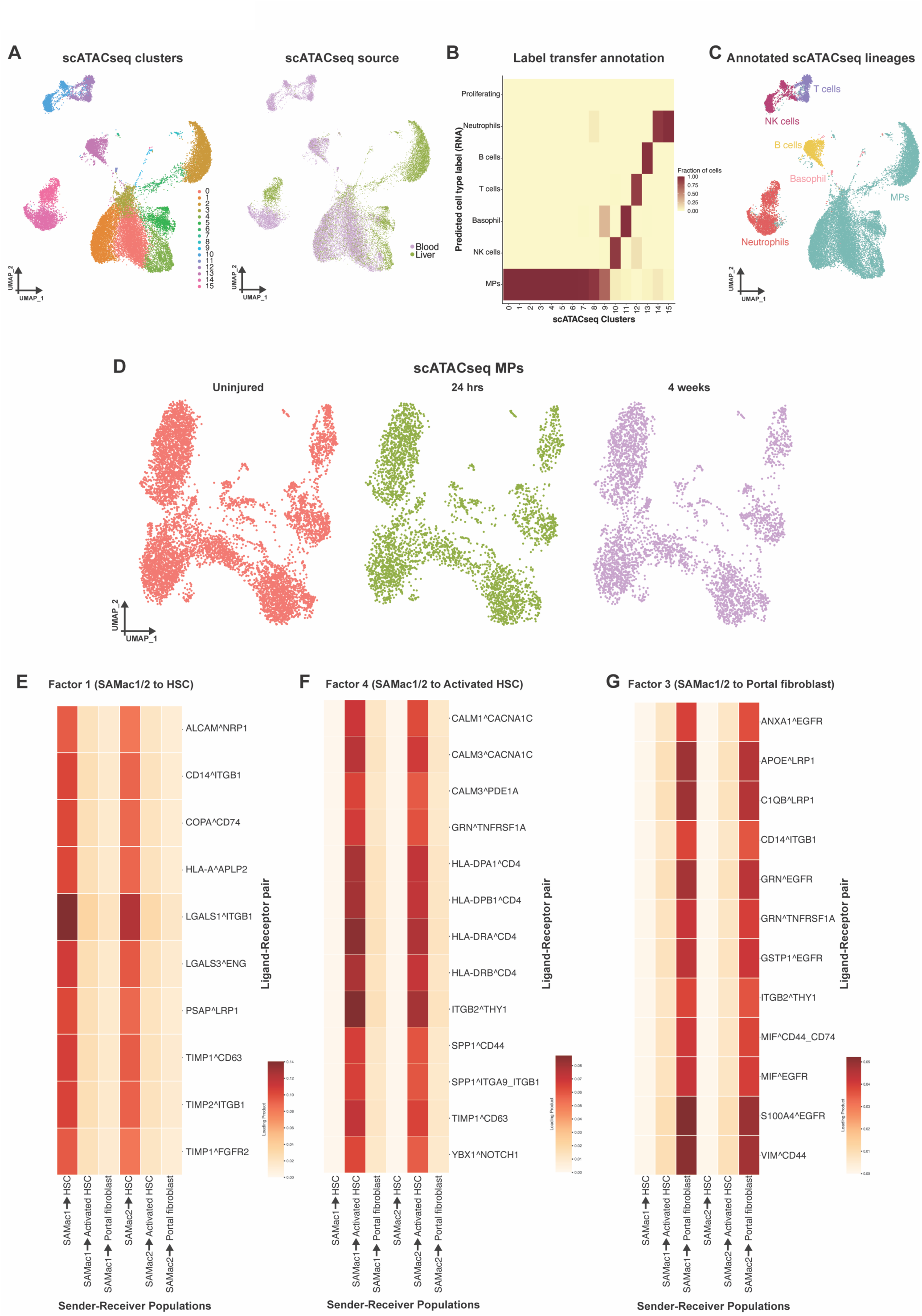
Identifying mouse MPs for subsequent scATACseq analysis. (**A**) UMAPs of mouse immune cell scATACseq data (27, 272 cells) from blood and liver of chronic CCl_4_ model (see Figure 5A). Coloured and labelled by immune cell scATACseq clusters (left) and by tissue source blood vs liver (right). (**B**) Heatmap of label transfer annotation analysis between mouse immune cell scATACseq clusters and annotated mouse immune cell scRNAseq clusters (from Extended Data Fig. 8A). Colour scale indicates prediction score of annotation of each scATACseq cluster. The highest prediction scores were used to annotate immune cell scATACseq clusters. (**C**) UMAP of mouse immune cell scATACseq data coloured and labelled by cell type annotation. MPs were then subsetted for further analysis. (**D**) UMAPs of mouse MP scATACseq data (see Figure 5B), coloured and split by condition/CCl_4_ model timepoint. (**E**-**G**) Supplementary heatmaps of ligand-receptor interaction analysis in human level 3 atlas shown in Figure 5I. Each factor represents a different cell-cell communication program. (**E**) Heatmap of selected ligand-receptor pairs in factor 1. Rows represent ligand-receptor pairs; columns represent specific cell type interactions; colour scale represents the loading product (degree of enrichment). (**F**) Heatmap of selected ligand-receptor pairs in factor 4. Rows represent ligand-receptor pairs; columns represent specific cell type interactions; colour scale represents the loading product (degree of enrichment). (**G**) Heatmap of selected ligand-receptor pairs in factor 3. Rows represent ligand-receptor pairs; columns represent specific cell type interactions; colour scale represents the loading product (degree of enrichment).

**Extended Data Fig. 11:**
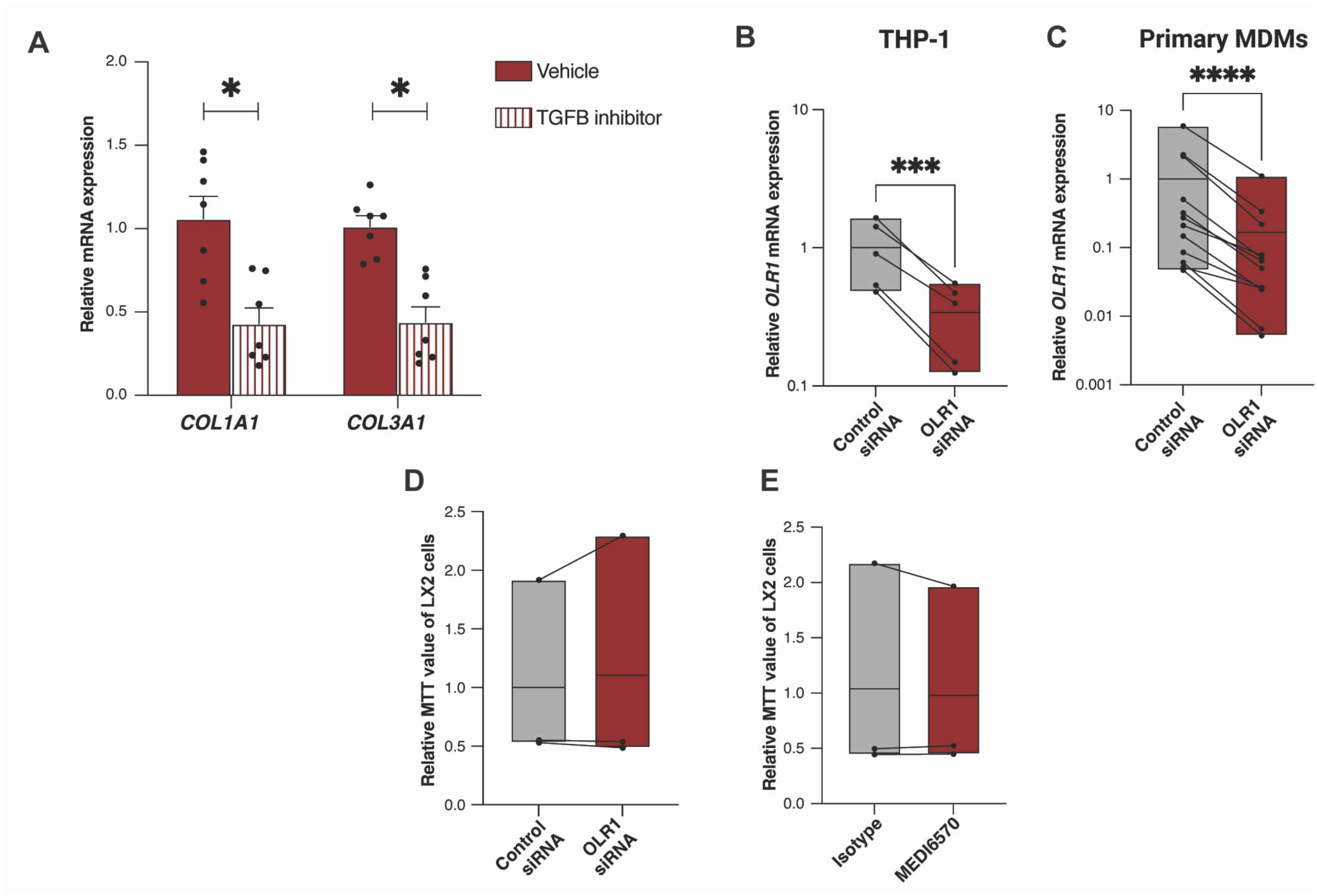
Modulation of macrophage OLR1 *in vitro*. (**A**) qPCR of D15 human liver spheroids (see Figure 6A) comparing 7 days treatment with TGFβ inhibitor (A83-01) or vehicle control. Dots represent biological replicates with monocytes from 7 different CLD patients. Expression of *COL1A1* and *COL3A1* in spheroids relative to mean expression of control (vehicle) group is shown. Bars indicate mean ± S.E.M. for each group. Wilcoxon matched-pairs signed rank test performed between groups for each gene; \**p*<0.05. (**B**) Relative *OLR1* mRNA expression in differentiated THP-1 cells following treatment with OLR1 targeting or control siRNA. Bars are min to max expression with line at mean. Dots connected by lines represent data from 5 independent experiments. Ratio paired t-test, \*\*\**p*<0.001. (**C**) Relative *OLR1* mRNA expression of primary MDM following treatment with OLR1 targeting or control siRNA. Bars are min to max expression with line at mean. Dots connected by lines represent data from 12 different CLD patients. Wilcoxon matched-pairs signed rank test, \*\*\**p*<0.001. (**D**) Relative MTT assay value of LX2 hepatic stellate cells following incubation with conditioned media from primary MDMs pre-treated with OLR1 targeting or control siRNA. Bars are min to max MTT value with line at mean. Dots connected by lines represent data from MDMs of 3 different CLD patients. (**E**) Relative MTT assay value of LX2 hepatic stellate cells following incubation with conditioned media from primary MDMs pre-treated with OLR1 blocking antibody (MEDI6570) or isotype control antibody. Bars are min to max MTT value with line at mean. Dots connected by lines represent data from MDMs of 3 different CLD patients.

## Supplemental Information

**Supplementary Table 1:** Summary of metadata for patients included in human scRNAseq analysis.

**Supplementary Table 2:** Level 1 cluster marker genes from human scRNAseq analysis.

**Supplementary Table 3:** Level 2 cluster marker genes from human scRNAseq analysis.

**Supplementary Table 4:** Level 3 cluster marker genes from human scRNAseq analysis.

**Supplementary Table 5:** Level 3 mononuclear phagocyte (MP) marker genes from human scRNAseq analysis.

**Supplementary Table 6:** Level 3 mesenchyme marker genes from human scRNAseq analysis.

**Supplementary Table 7:** Level 3 endothelia marker genes from human scRNAseq analysis.

**Supplementary Table 8:** Level 3 T/NK cell marker genes from human scRNAseq analysis.

**Supplementary Table 9:** Level 3 B cell marker genes from human scRNAseq analysis.

**Supplementary Table 10:** Level 3 epithelial marker genes from human scRNAseq analysis.

**Supplementary Table 11:** Differentially expressed genes between human SAMac1 and SAMac2.

**Supplementary Table 12:** Cluster marker genes for mouse mononuclear phagocytes.

**Supplementary Table 13:** Differentially expressed genes between mouse SAMac1 and SAMac2.

**Supplementary Table 14:** Transcription factor motif enrichment in mouse SAMac1 (tab 1) and SAMac2 (tab 2) from scATACseq data.

**Supplementary Table 15:** Ligand-receptor interaction analysis from human scRNAseq data between SAMac1 and SAMac2 as senders and level 3 mesenchyme as receiver cells. Summary of ligand-receptor pairs in each factor.

**Supplementary Table 16:** Antibodies used in this study.

**Supplementary Table 17:** qPCR primers used in this study.

